# Assessing the Capacity of Genomic Language Models to Generate Informative Embeddings of Viral Genome Fragments

**DOI:** 10.64898/2026.11.09.765411

**Authors:** A. Kladchenko, N.S. Popov, A.I. Manolov

## Abstract

Genomic language models (gLMs) trained on large volumes of nucleotide sequences are becoming an alternative to classical methods in metagenomic analysis, particularly for studying genomic “dark matter”. In this work, we investigate the ability of four gLMs (DNABERT-2, DNABERT-S, ViraLM, and Evo2-7B) to generate taxonomically informative vector representations of short viral genome fragments. We constructed a balanced reference dataset of viruses based on the ICTV Virus Metadata Resource and compared the representations generated by gLMs with conventional k-mer profiles and MinHash sketches using clustering and neighborhood-based metrics (adjusted Rand index (ARI), precision@k, and CROSS precision@k).

On the reference dataset, three of the four gLMs outperformed the 4-mer profile in terms of ARI and precision@5 under the same evaluation protocol. The best performance was generally achieved by intermediate model layers. Conventional k-mer profiles consistently outperformed MinHash sketches. We further evaluated the methods on two viral orders (egoviruses and mriyaviruses) absent from the reference dataset, with model layers and k-mer lengths selected exclusively on the reference dataset. DNABERT-S showed the most consistently strong performance across these two orders, ranking first on mriyaviruses and second on egoviruses, while k-mer profiles with Jensen–Shannon distance remained competitive. We also observed the consistency of representations between forward fragments and their reverse complements.

Overall, the best-performing gLMs surpassed conventional baselines, although their advantage depended on the dataset and evaluation metric. The consistently strong performance of DNABERT-S suggests a potential benefit of contrastive fine-tuning and indicates that larger models are not necessarily more effective for generating taxonomically informative representations.

## 1 Introduction

Viruses are important both for ecosystems and for public health; however, the vast majority of them cannot be cultured, so metagenomic sequencing has become the primary tool for studying viral diversity. At the same time, the analysis of viral metagenomes is inherently complex: viruses are extremely diverse, evolve rapidly, and lack universal marker genes. The data usually consist of short fragments (contigs) [1].

Traditional metagenomic workflows rely on sequence alignment and k-mer-based methods to classify or cluster sequence fragments. Alignment-based classification depends on the availability of homologous reference sequences, while k-mer-based methods primarily capture local sequence composition and do not explicitly account for long-range dependencies. Although k-mer-based approaches can be used without reference databases, assigning taxonomic identities to highly divergent or previously uncharacterized sequences remains challenging. Consequently, a substantial fraction of metagenomic sequences remains unclassified, forming what is commonly referred to as metagenomic “dark matter”.

These limitations motivate the search for approaches that do not rely directly on reference database coverage. One important task in the analysis of metagenomic “dark matter” is to identify groups of sequence fragments that originate from the same genome, an approach known as metagenomic binning. Such groups can then be studied across samples to investigate their distribution and abundance. For example, sequences associated with a particular ecological niche may provide clues to the presence of previously uncharacterized viruses. Methods that capture biologically meaningful similarities between sequences, even in the absence of known homologs, may therefore be useful for the exploration of viral diversity.

Language models trained on genomic data (genomic language models, gLMs) can extract hidden functional and evolutionary structure directly from nucleotide sequences [2]. A transformer-based deep-learning model of protein sequences has already made it possible to describe a previously unknown part of the virosphere, identifying more than 1.6 × 10^5^ putative new species of RNA viruses [3]. Despite these advances, it remains unclear how well gLMs capture taxonomically informative features of short viral genome fragments and whether their representations outperform conventional sequence-based methods. It is also uncertain how representation quality varies between models and across their layers. In this study, we address these questions using a balanced reference dataset and two additional viral orders excluded from it. We compare four gLMs with three conventional baselines, selecting model layers and k-mer lengths exclusively on the reference dataset to assess how well the resulting representations generalize to other viruses.

## 2 Materials and Methods

### 2.1 Reference dataset and class balancing

To construct a reference dataset, we used the expert-curated ICTV Virus Metadata Resource (VMR, October 13, 2025) [4]. To reduce class imbalance, we applied the following sampling procedure:

1. Taxon selection We retained viral classes containing more than 10 species.
2. Genome selection and fragmentation For each class, we randomly selected 10 genomes and divided them into non-overlapping fragments of 2,000 nucleotides.
3. Fragment sampling From each genome, we randomly selected up to 10 fragments.

We repeated steps 2–3 50 times, generating 50 random subsamples. Each subsample contained fragments from 39 viral classes. On average, each subsample contained about 1,300 fragments, corresponding to about 33 fragments per class. The number of fragments varied between classes because shorter genomes yielded fewer complete 2,000-nucleotide fragments. Repeated sampling allowed us to assess the variability of the evaluation metrics arising from the random selection of genomes and fragments.

#### 2.2 Held-out viruses

To evaluate the representations on viruses outside the reference dataset, we selected two viral orders not included in it: egoviruses [5] and mriyaviruses [6]. We processed their genomes using the same fragmentation and sampling procedure as for the reference dataset (see Limitations for details on the dataset version used in the analysis). We also generated reverse complements of the fragments to assess the consistency of the representations between the two sequence orientations. We did not determine whether sequences from these viral orders were included in the models’ pretraining datasets.

### 2.3 gLM models

We evaluated four genomic language models (DNABERT-2 [7], DNABERT-S [8], ViraLM [1], and Evo2-7B [9]) and compared their performance with three conventional baselines: a 4-mer frequency profile with cosine distance, a k-mer frequency profile with Jensen–Shannon distance [10, 11], and MinHash sketches [12, 13] (Table 1). DNABERT-2 and DNABERT-S belong to the DNABERT family [14] of transformer-based models [15].

**Table 1.** Overview of the methods compared in this study.

| **Method** | **Representation / architecture** | **Sequence encoding** | **Layers** |
|---|---|---|---|
| 4-mer spectrum | frequency vector, cosine distance | overlapping k-mers | – |
| k-mer profile (JS) | frequency vector (4<sup>*k*</sup>, k = 4),<br>Jensen–Shannon distance | overlapping k-mers | – |
| MinHash | 128-hash MinHash sketch (k = 6) | canonical k-mers | – |
| DNABERT-2 (117M) | transformer encoder | BPE | 12 |
| DNABERT-S | transformer encoder (contrastive) | BPE | 12 |
| ViraLM | transformer encoder (fine-tuned) | BPE | 12 |
| Evo2-7B | StripedHyena 2 | single nucleotide | 32 |

### 2.4 Sequence representations

For DNABERT-2, DNABERT-S, and ViraLM, we extracted hidden states from the output of each encoder block. For Evo2-7B, we used the output of the feed-forward sublayer of each block (module mlp.l3, before the residual connection). The resulting hidden states formed a tensor of shape (1, *n_tok, d*), where *n_tok* is the number of token positions and *d* is the hidden-state dimension. We then applied mean pooling across the token dimension to obtain a d-dimensional vector representation for each layer.

For DNABERT-2 and DNABERT-S, mean pooling was performed without a padding mask, so padding positions were included in the average, at least at the embedding layer (see Limitations). For ViraLM, each 1,000-nucleotide window produced a separate vector representation. Thus, the two windows of each fragment were treated as separate points in the representation space, both retaining the label of the original fragment (see Limitations).

Layer numbering differed between models. For DNABERT-2 and DNABERT-S, layer 0 corresponds to the embedding output, and layers 1–12 correspond to the encoder blocks. For ViraLM, layers 1–12 correspond to its encoder blocks, with the embedding output excluded from the analysis. For Evo2-7B, layers 0–31 correspond to its 32 blocks. This procedure yielded a set of layer-wise representations for each fragment, or for each window in the case of ViraLM (Figure 1).

**Figure 1.**
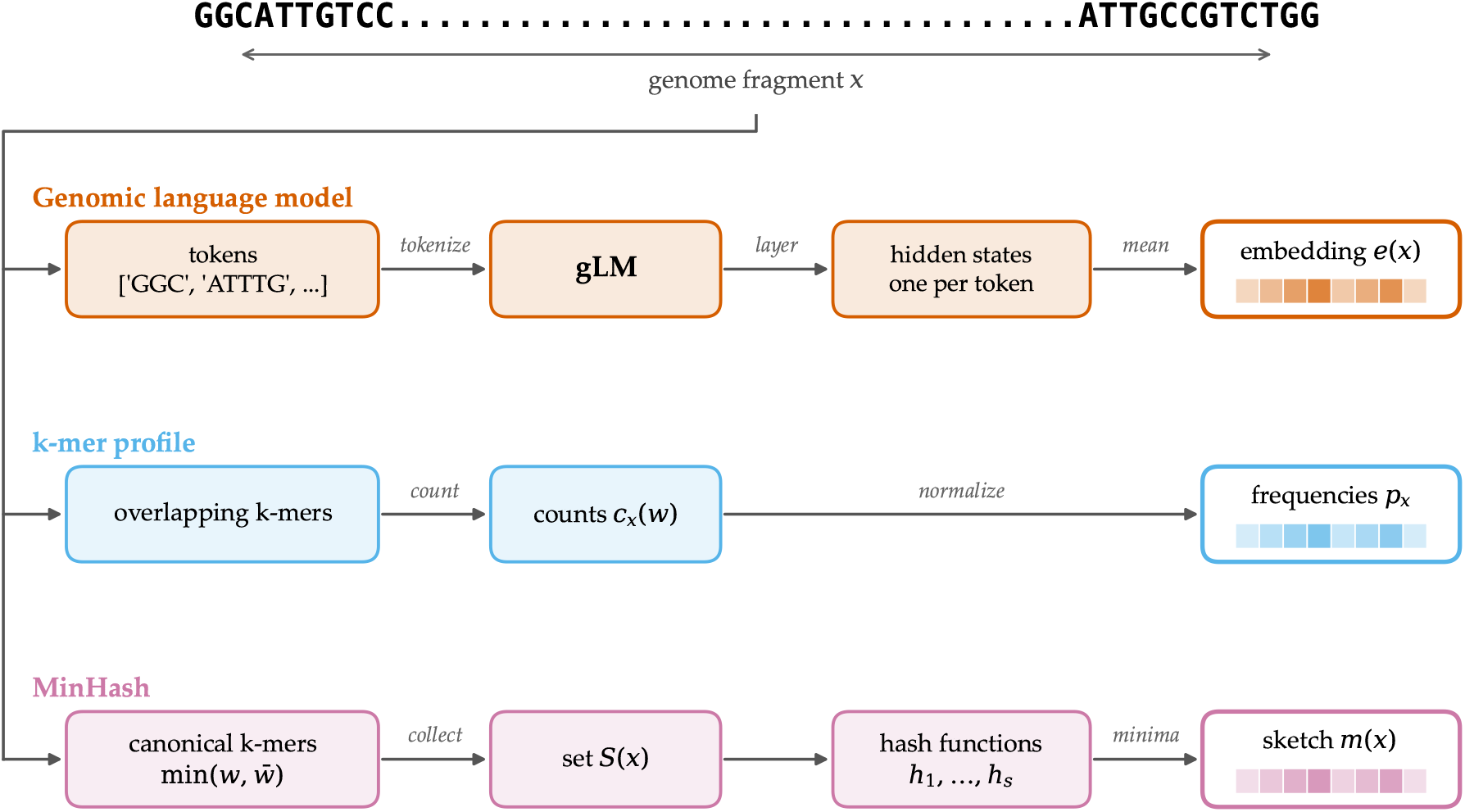
Overview of the sequence representations compared in this study. Each 2,000-nucleotide fragment is represented using three approaches. Genomic language models generate layer-wise hidden states, which are averaged across token positions to obtain vector representations. These vectors are standardized and compared using cosine distance. k-mer frequency profiles are compared using either cosine distance after standardization or Jensen–Shannon divergence without standardization. MinHash represents each fragment as a set of canonical k-mers summarized by a sketch of 128 hash values. The distance between two sketches is calculated as the fraction of positions at which they differ.

For each fragment, we constructed a k-mer frequency profile by counting all overlapping nucleotide sequences of length k. Only k-mers consisting of the four standard nucleotides (A, C, G, and T) were considered; windows containing other characters (e.g., N) were excluded. The counts were then normalized by the total number of valid k-mers in the fragment, yielding a frequency vector of length 4*^k^* whose elements sum to one. The optimal value of k was selected using the reference dataset, as described in Section 2.5.

### 2.5 Parameter selection

For each gLM, we selected the model layer that maximized mean precision@5 across the 50 reference subsamples. Similarly, the k-mer length for each conventional baseline was selected by maximizing mean precision@5, using 10 reference subsamples. We tested k-mer lengths from 2 to 8 for frequency profiles and from 2 to 10 for MinHash. The optimal length was k = 4 for both k-mer frequency profile methods and k = 6 for MinHash.

All parameters were selected exclusively on the reference dataset and kept fixed for the evaluation of egoviruses and mriyaviruses. Thus, the results on these viral orders were obtained without optimizing parameters on the held-out data.

### 2.6 Evaluation metrics

To evaluate how well the different methods capture taxonomic relatedness between viral sequences, we used the following clustering and neighborhood-based metrics.

#### Precision@k (P@k)

measures the average proportion of the k nearest neighbors that belong to the same class as the query fragment. Neighbors were identified using the distance measure associated with each method. In the reference dataset, fragments were labeled according to their ICTV taxonomic class. Other fragments of the same genome were not excluded from the neighborhood, so the metric reflects both similarity between fragments of the same genome and similarity between different genomes of the same class.

#### Adjusted Rand index (ARI)

[16] measures the agreement between the true class labels and the clustering obtained using k-medoids [17], corrected for chance. The number of clusters was set equal to the number of taxonomic classes. For the Jensen–Shannon and MinHash baselines, we used our own implementation of k-medoids. For the other methods, we used the KMedoids implementation from scikit-learn-extra.

#### Silhouette coefficient

[18] was calculated using the true class labels. Since its values depend on the distance measure and are not directly comparable across methods using different distances, the results are reported only in the Supplementary Materials.

#### CROSS precision@k (CROSS P@k)

measures the proportion of the k nearest neighbors of a query fragment that belong to a specified target set. We used this metric in two settings:

1. **Reverse-complement (RC) proximity** For each forward fragment, we measured the proportion of its nearest neighbors that were reverse-complement fragments from the same genome. The search space consisted of one reference subsample and the reverse complements of the fragments of the corresponding genome, excluding the original forward fragments. A neighbor was counted if it was the reverse complement of any fragment from the same genome, not necessarily of the query fragment itself.
2. **Inter-order overlap** We measured the proportion of mriyaviruses fragments among the nearest neighbors of egoviruses fragments. The search space consisted of one reference subsample and the mriyaviruses fragments. Lower values indicate better separation between the two viral orders, while the value expected by chance corresponds to the proportion of mriyaviruses fragments in the search space.[1]

For the held-out viruses, precision@k was calculated using genome identity rather than taxonomic class as the label, with the search space consisting of fragments from the corresponding genome and one reference subsample. The metric thus measured the proximity of fragments from the same genome rather than taxonomic classification accuracy. Precision@k and RC proximity were first averaged across fragments within each genome and then across genomes, giving each genome equal weight. Inter-order overlap was averaged across all egoviruses fragments. All results were then averaged over the 50 reference subsamples. Where applicable, feature standardization was performed using the entire search space.

### 2.7 Statistical analysis

All methods were evaluated on the same 50 reference subsamples, allowing paired statistical comparisons. Differences between methods were assessed using the two-sided Wilcoxon signed-rank test [19], applied to the metric values obtained for each subsample. To account for multiple testing, p-values were adjusted using the Benjamini–Hochberg procedure [20]. For the held-out viruses, the correction was applied across504 pairwise comparisons (21 pairs of methods across 24 evaluation conditions; Sections 3.2 and 3.3). Other comparisons, including those between individual methods and the 4-mer baseline, between model layers or k-mer lengths, were corrected separately within each figure (Sections 3.1 and 3.1.1). Exact p-values were calculated where possible; a normal approximation was used when tied differences prevented exact calculation.

Importantly, the 50 reference subsamples were drawn from the same pool of genomes and therefore overlapped. Moreover, the held-out genomes remained unchanged across evaluations, with only the reference background varying. Consequently, these statistical comparisons assess the robustness of the observed differences to reference subsampling rather than their generalizability to other viral genomes.

## 3 Results

### 3.1 Reference benchmark

First, we measured how well each method clusters fragments of the same reference class using precision@5 (P@5) to measure local neighborhood homogeneity and the adjusted Rand index (ARI) to assess global clustering performance (Figure 2, Table 2).

**Table 2.**
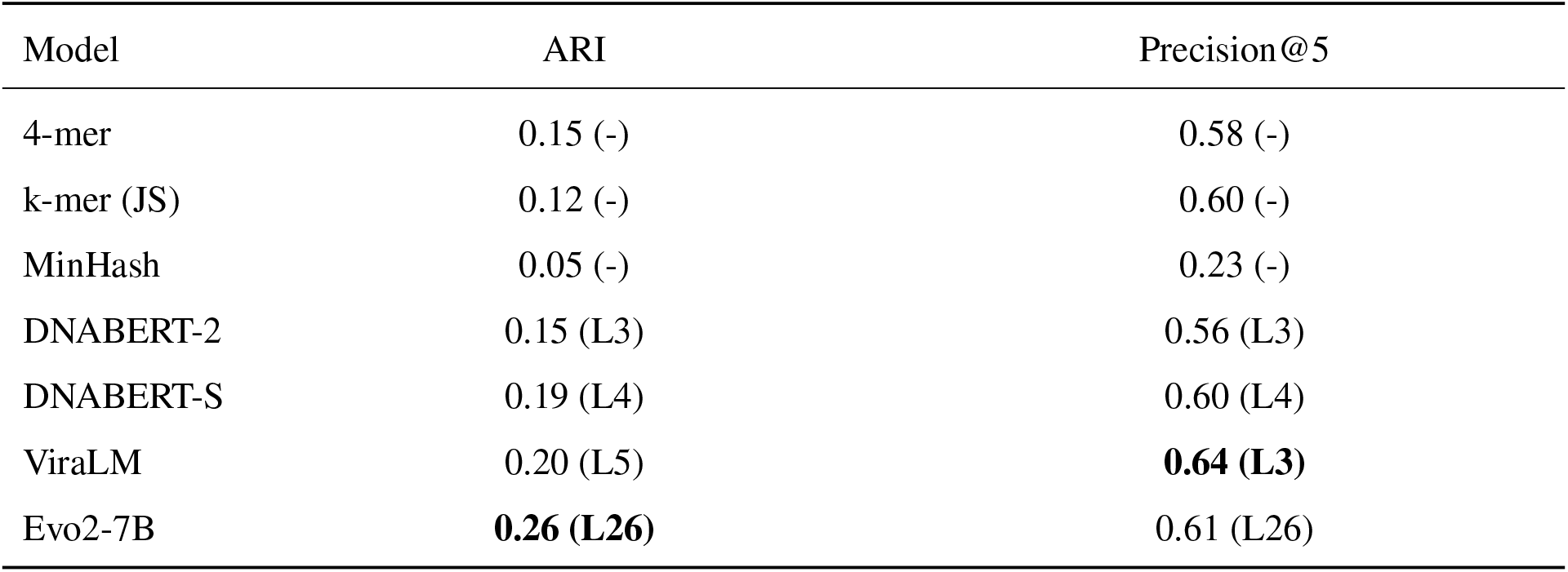
Best layer-wise scores on the reference set; each metric is reported at its own optimal layer (layer index in parentheses). Layers are numbered from 0.

| Model | ARI | Precision@5 |
|---|---|---|
| 4-mer | 0.15 (-) | 0.58 (-) |
| k-mer (JS) | 0.12 (-) | 0.60 (-) |
| MinHash | 0.05 (-) | 0.23 (-) |
| DNABERT-2 | 0.15 (L3) | 0.56 (L3) |
| DNABERT-S | 0.19 (L4) | 0.60 (L4) |
| ViraLM | 0.20 (L5) | **0.64 (L3)** |
| Evo2-7B | **0.26 (L26)** | 0.61 (L26) |

**Figure 2.**
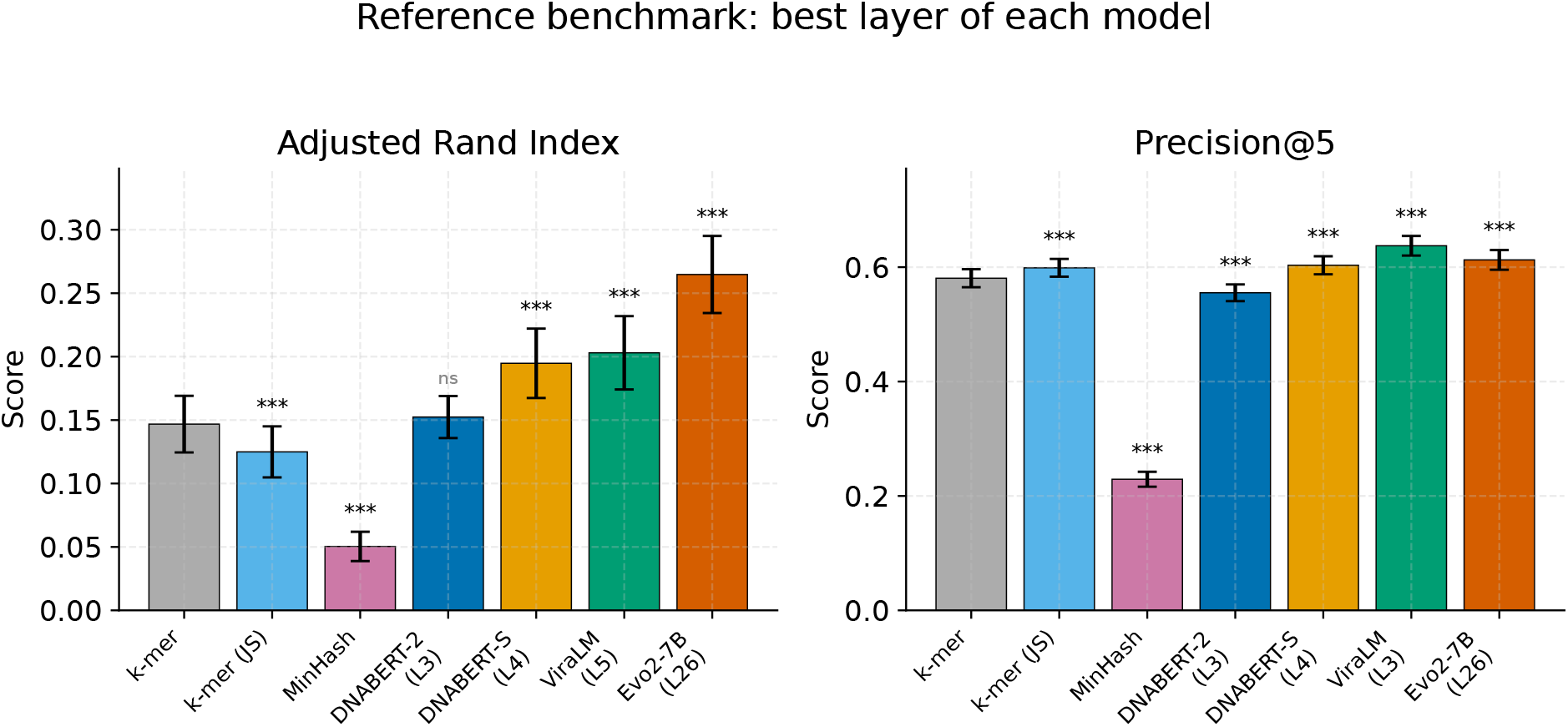
Performance of genomic language models and conventional methods on 50 reference subsamples from the ICTV VMR dataset. Adjusted Rand index (ARI) and precision@5 are shown as mean ± standard deviation. For each gLM, the optimal layer was selected separately for each metric, with layer indices indicated above the bars. Statistical significance is shown relative to the k-mer frequency profile baseline. Asterisks show Benjamini–Hochberg-adjusted p-values of two-sided paired Wilcoxon tests over the reference subsets: * p < 0.05, ** p < 0.01, *** p < 0.001; ns, not significant. Each method is compared with the 4-mer profile.

Three of the four gLMs (ViraLM, DNABERT-S, and Evo2-7B) outperformed the conventional 4-mer frequency profile on both metrics, although the improvements in P@5 were modest (0.02–0.06). DNABERT-2 showed no improvement over the baseline (ARI: 0.15 vs. 0.15; P@5: 0.56 vs. 0.58). Among the gLMs, ViraLM achieved the highest P@5 (0.64), while Evo2-7B showed the highest ARI (0.26). However, the ViraLM results may be inflated by the treatment of its 1,000-nucleotide windows as separate points (see Limitations).

Among the conventional methods, the k-mer frequency profile with Jensen–Shannon distance achieved a P@5 of 0.60, matching DNABERT-S and exceeding DNABERT-2. However, its ARI (0.12) was lower than that of the 4-mer profile with cosine distance (0.15). MinHash performed substantially worse than all other methods, with a P@5 of 0.23 and an ARI of 0.05. Both alternative baselines differed significantly from the 4-mer profile on both metrics (Figure 2).

For the gLMs, the highest scores were generally obtained from intermediate rather than final layers (Figure 3, Table 2). Evo2-7B also showed a secondary local maximum at layer 3, which is examined further in Sections 3.4 and 4. No single model achieved the highest performance on both metrics.

**Figure 3.**
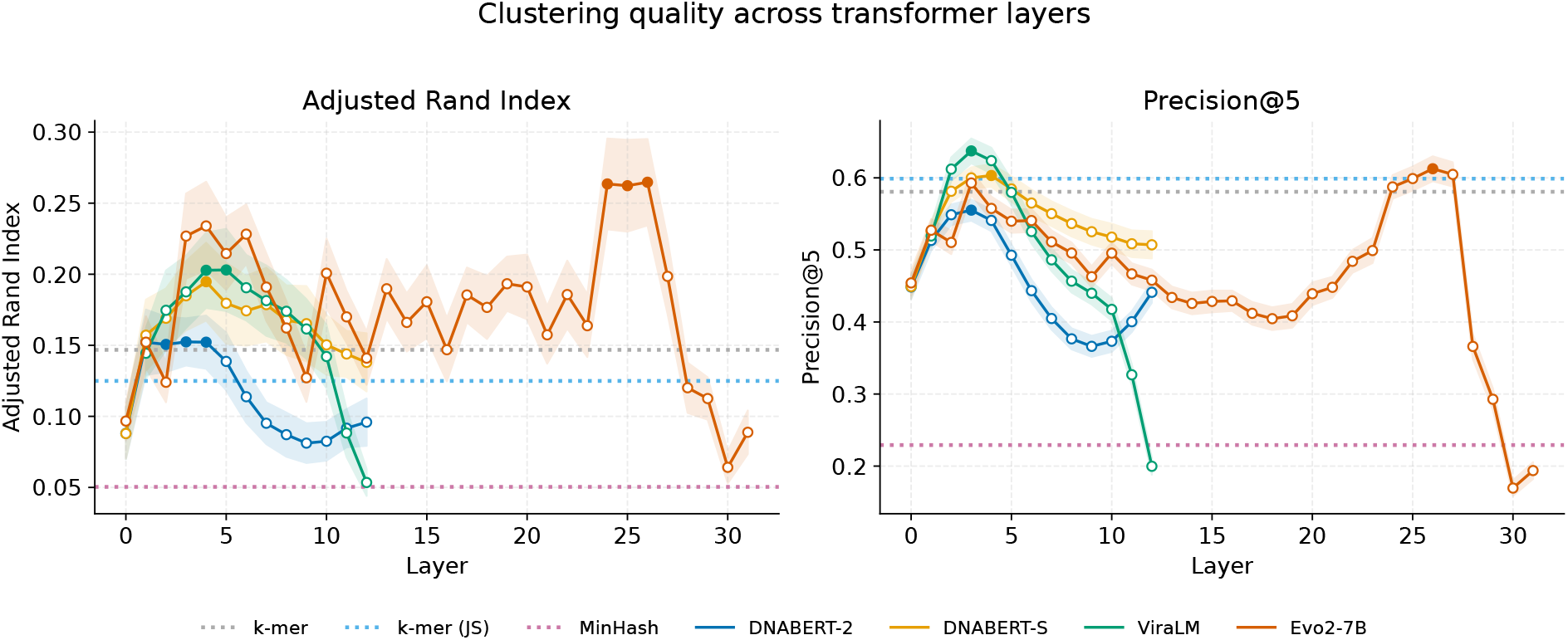
Layer-wise ARI and precision@5 for each model (mean across subsamples; dotted lines, classical baselines: k-mer, k-mer (JS), MinHash).

**Figure 4.**
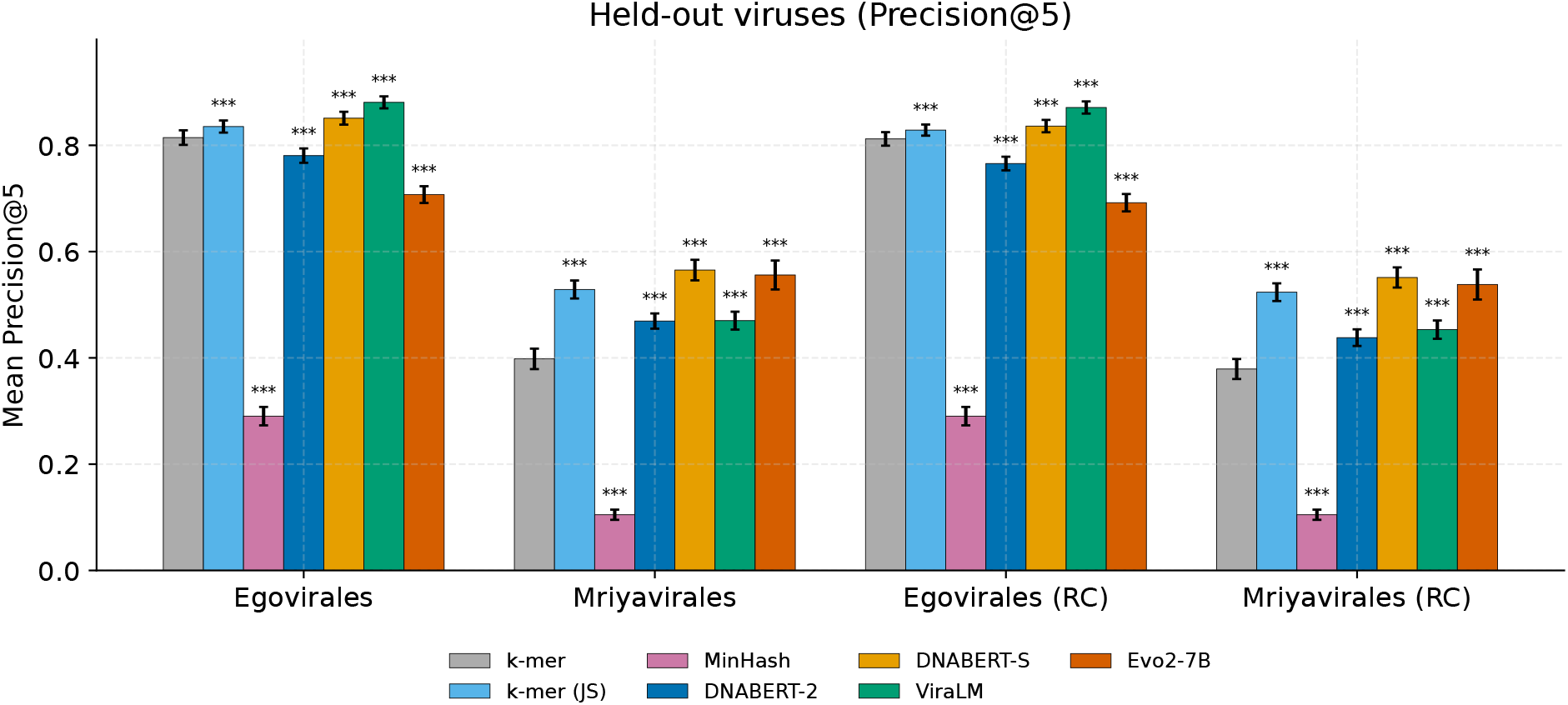
Held-out viruses: precision@5 for each model on egoviruses and mriyaviruses and their reverse complements (RC). Mean ± standard deviation across 50 subsets. The layer of each model is chosen on the reference set. Asterisks show Benjamini–Hochberg-adjusted p-values of two-sided paired Wilcoxon tests over the reference subsets: * p < 0.05, ** p < 0.01, *** p < 0.001; ns, not significant. Each method is compared with the 4-mer profile.

#### 3.1.1 choosing k for the k-mer baselines

A fair comparison with the classical descriptors requires their best-performing k-mer length, which was chosen by P@5 on the reference subsets (Figure S1). For the k-mer profile with the cosine distance both ARI and P@5 peaked at k = 4 (P@5 = 0.58), and for the k-mer profile with the JS distance the best value was also k = 4 (P@5 = 0.61 over 10 subsets; Table 2 gives the mean over all 50 subsets). MinHash was run with k from 2 to 10; its P@5 was highest at k = 6 (0.23), at the edge of this range, and decreased as k grew.

### 3.2 Performance on held-out viruses

To evaluate the representations on viruses outside the reference dataset, we selected two recently described groups of viruses: egoviruses [5] and mriyaviruses [6]. We processed their genomes using the same fragmentation and sampling procedure as for the reference dataset.

For the held-out viruses, genome identity was used as the label, so P@5 measured the proximity of fragments from the same genome rather than their similarity to related genomes. The maximum possible P@5 depended on the number of fragments available from each genome.

On egoviruses, ViraLM (0.88) and DNABERT-S (0.85) outperformed the 4-mer baseline (0.81), while DNABERT-2 (0.78) and Evo2-7B (0.71) performed worse. On mriyaviruses, all four gLMs outperformed the baseline (0.40), with DNABERT-S (0.57) and Evo2-7B (0.56) achieving the highest scores. The k-mer frequency profile with Jensen–Shannon distance achieved P@5 values of 0.84 on egoviruses and 0.53 on mriyaviruses, outperforming DNABERT-2 and Evo2-7B on egoviruses and DNABERT-2 and ViraLM on mriyaviruses. MinHash performed substantially worse than all other methods (0.29 and 0.10, respectively). Overall, DNABERT-S showed the most consistently strong performance across both groups of held-out viruses, although ViraLM achieved the highest P@5 on egoviruses. The k-mer profile with Jensen–Shannon distance remained competitive with the gLMs, indicating that their advantage over conventional methods depends on the model and the viral dataset.

### 3.3 Forward fragments and their reverse complements

The P@5 values for forward fragments and their reverse complements were similar across all methods (Table 3). However, this agreement does not necessarily indicate that the models generate similar representations for the two sequence orientations, as comparable results were also observed for the conventional k-mer baselines. To examine this further, we evaluated reverse-complement (RC) proximity using CROSS P@5 (Figure 5, Table 4). For each forward fragment, this metric measured the proportion of its five nearest neighbors that were reverse complements of fragments from the same genome. Since any fragment from the same genome counted as a match, the metric reflects genome-level proximity rather than direct similarity between a fragment and its own reverse complement.

**Table 3.** Precision@5 for the held-out viruses (layer chosen on the reference set; average across subsets).

| Model | egoviruses | mriyaviruses | egoviruses RC | mriyaviruses RC |
|---|---|---|---|---|
| k-mer | 0.81 | 0.40 | 0.81 | 0.38 |
| k-mer (JS) | 0.84 | 0.53 | 0.83 | 0.52 |
| MinHash | 0.29 | 0.10 | 0.29 | 0.10 |
| DNABERT-2 | 0.78 | 0.47 | 0.77 | 0.44 |
| DNABERT-S | 0.85 | **0.57** | 0.84 | **0.55** |
| ViraLM | **0.88** | 0.47 | **0.87** | 0.45 |
| Evo2-7B | 0.71 | 0.56 | 0.69 | 0.54 |

**Table 4.** Reverse-complement (RC) proximity (CROSS precision@5) for egoviruses and mriyaviruses.

| Model | egoviruses | mriyaviruses |
|---|---|---|
| k-mer | 0.61 | 0.34 |
| k-mer (JS) | 0.65 | 0.49 |
| MinHash | 0.45 | 0.29 |
| DNABERT-2 | 0.65 | 0.47 |
| DNABERT-S | 0.77 | **0.59** |
| ViraLM | **0.78** | 0.46 |
| Evo2-7B | 0.52 | 0.58 |

**Figure 5.**
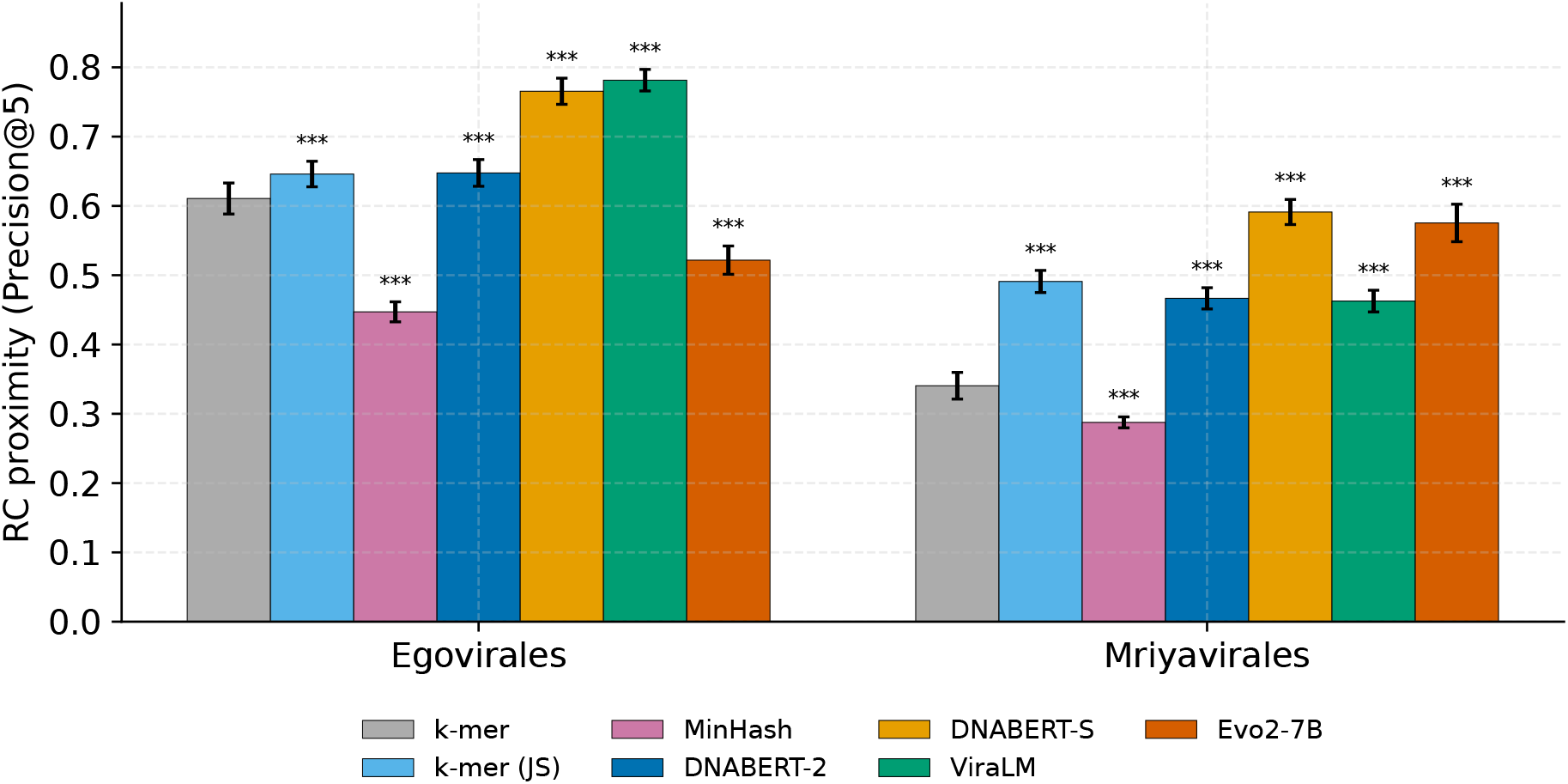
Reverse-complement (RC) proximity (CROSS precision@5) for egoviruses and mriyaviruses. Values are shown as mean ± standard deviation across 50 reference subsamples. Each method was compared with the 4-mer frequency profile using a paired two-sided Wilcoxon signed-rank test, with p-values adjusted for multiple comparisons using the Benjamini–Hochberg procedure, each method is compared with the 4-mer profile. Statistical significance is indicated as follows: * p < 0.05, ** p < 0.01, *** p < 0.001.

On egoviruses, ViraLM (0.78) and DNABERT-S (0.77) achieved the highest RC proximity, while DNABERT-S (0.59) and Evo2-7B (0.58) performed best on mriyaviruses. Among the gLMs, only Evo2-7B on egoviruses (0.52) performed worse than the 4-mer baseline (0.61 on egoviruses and 0.34 on mriyaviruses).

MinHash showed considerably lower values (0.45 and 0.29, respectively), despite using canonical k-mers that produce identical sketches for a fragment and its reverse complement. This indicates that its low RC proximity reflects poor separation of fragments from different genomes rather than sensitivity to sequence orientation.

The relative performance of the methods remained largely stable across different neighborhood sizes (Figure 6). On mriyaviruses, Evo2-7B achieved the highest P@k for k = 1–3, while DNABERT-S performed best at larger k. ViraLM showed the smallest decline in P@k as the neighborhood size increased, although this result may be influenced by its representation of each fragment as two separate points, increasing the number of potential same-genome neighbors (Section 2.4).

**Figure 6.**
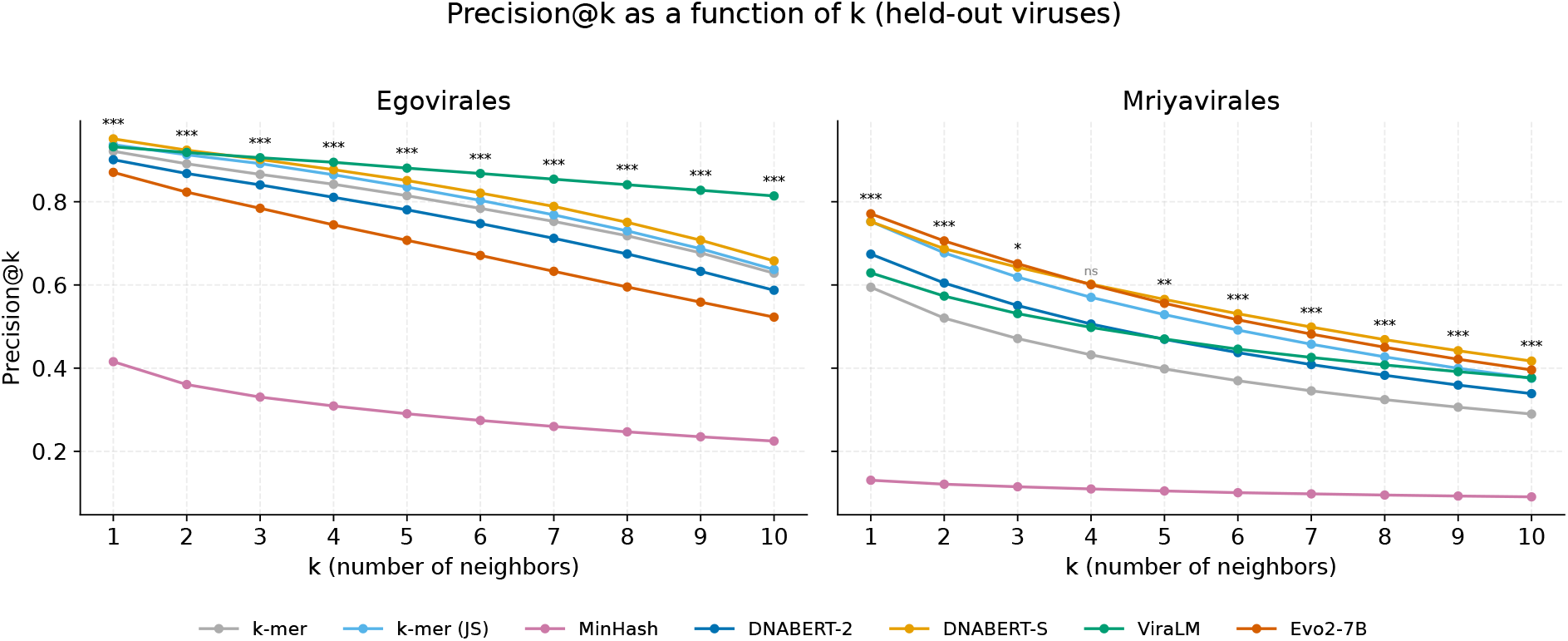
Dependence of precision@k on the neighborhood size k for the held-out egoviruses and mriyaviruses. Asterisks show Benjamini–Hochberg-adjusted p-values of two-sided paired Wilcoxon tests over the reference subsets: * p < 0.05, ** p < 0.01, *** p < 0.001; ns, not significant. At each k, the leading method is compared with the second one.

RC proximity: share of the 5 nearest neighbors that are reverse-complement fragments of the SAME genome (not necessarily the fragment’s own RC)

#### 3.4 Do the held-out viruses cluster into a single region?

If the input at inference differs substantially from the training data, the model may produce unstable representations. In particular, fragments that differ strongly from the training sequences may cluster in a single region of the representation space. To examine this possibility, we plotted intra-genome proximity (P@10) against inter-group overlap (CROSS P@10) for each layer (Figure 7). Here, CROSS P@10 is the proportion of mriyavirus fragments among the 10 nearest neighbors of an egovirus fragment when the search space consists of one reference subsample and the mriyavirus fragments; lower values indicate better separation. P@10 is averaged across the two viral groups, each evaluated with the other group in the search space. The ideal representation lies in the upper-left corner, indicating high intra-genome proximity and low inter-group overlap. ViraLM layers 2 and 3 and Evo2-7B layer 3 lie in this region, and all DNABERT-S layers have CROSS P@10 below 0.24; this suggests that the models do not place all held-out fragments into a single undifferentiated cluster. The 4-mer baseline also lies in this region: it has the lowest CROSS P@10 of all points (0.12) and a P@10 of 0.45, comparable to the best layer of DNABERT-2. The best-performing layers on these viruses differ from those selected on the reference set: Evo2-7B layer 26 has a P@10 of 0.43 and a CROSS P@10 of 0.26, performing worse than the 4-mer baseline on both axes.

**Figure 7.**
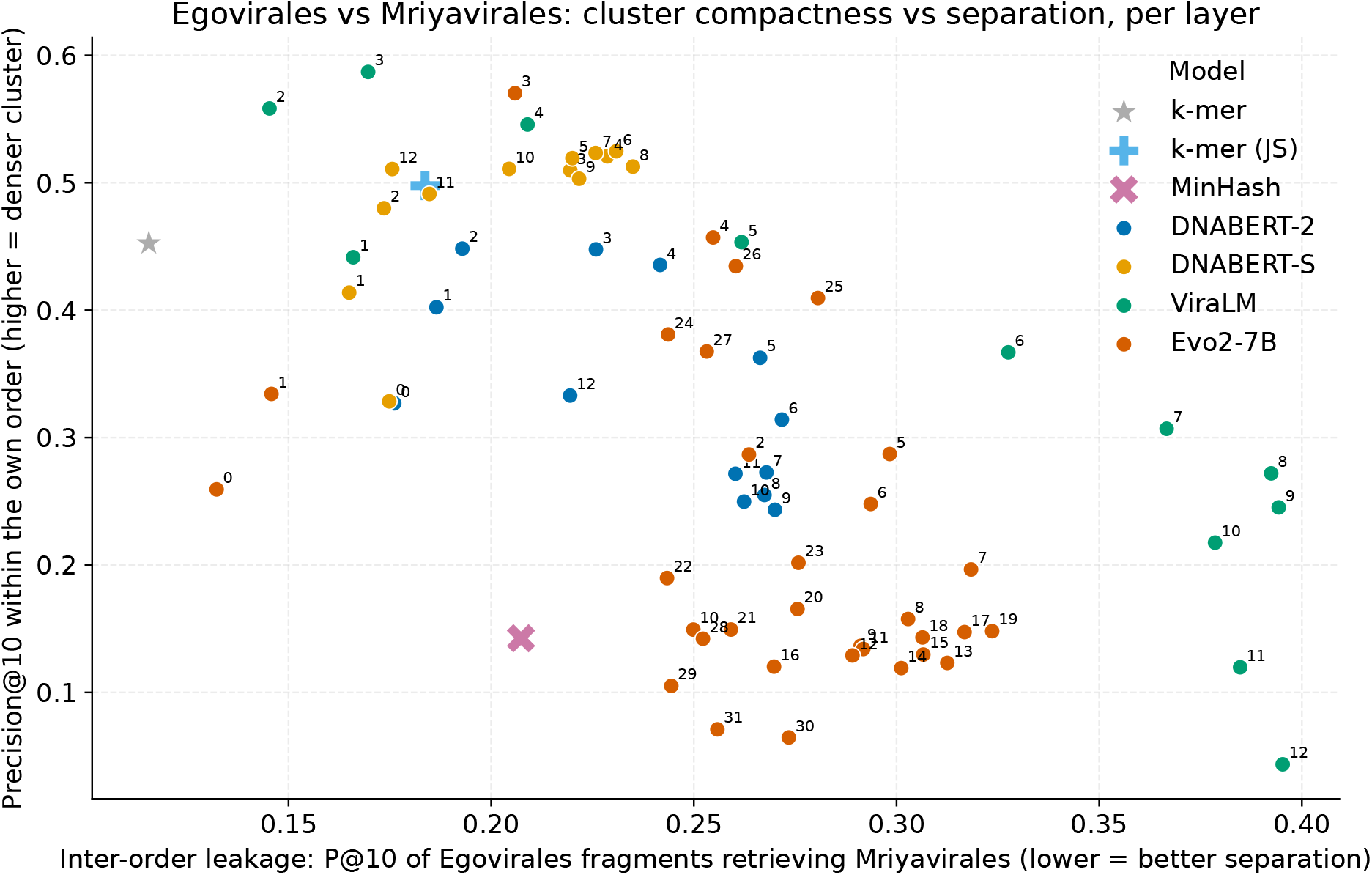
Layer-by-layer trade-off between intra-genome density (P@10) and inter-order overlap (CROSS P@10) for the held-out viruses; each point represents a layer, labeled with its number. The ideal region is in the upper left. The k-mer (JS) and MinHash representations are not included.

## 4 Discussion

On the class-balanced reference benchmark, three of the four genomic language models (ViraLM, DNABERT-S, and Evo2-7B) produced more taxonomically informative representations of short viral genome fragments than the conventional 4-mer frequency profile, as measured by ARI and P@5. However, the improvements were moderate, and DNABERT-2 did not outperform the baseline. The advantage of gLMs was also less pronounced when the k-mer profile was compared using Jensen–Shannon distance rather than standardized cosine distance. On the held-out egoviruses and mriyaviruses, performance varied considerably between models, with DNABERT-S showing the most consistently strong results.

An important finding is that the most informative representations were generally obtained from interme-diate model layers rather than the final layer. Moreover, the layers selected on the reference dataset were not always optimal for the held-out viruses. For Evo2-7B, layer 3 performed substantially better on the held-out viruses than layer 26, which achieved the highest score on the reference dataset. However, the alternative layer-selection strategy was identified after examining the results and requires further validation. No single model dominated across all metrics. ViraLM achieved the highest local homogeneity on the reference dataset and egoviruses, while Evo2-7B performed best in terms of ARI. DNABERT-S showed consistently strong performance across both held-out viral groups and in RC proximity. This suggests that contrastive fine-tuning may be beneficial for learning taxonomically informative representations, although its contribution cannot be isolated from other differences between the models. These results also indicate that larger models are not necessarily more effective for this task. The potential benefits of combining representations from different gLMs remain to be investigated.

The choice of distance measure substantially affected the performance of conventional methods. On mriyaviruses, the same 4-mer frequency profile achieved P@5 values of 0.40 with standardized cosine distance and 0.53 with Jensen–Shannon distance. On the reference dataset, Jensen–Shannon distance improved P@5 (0.60 vs. 0.58) but reduced ARI (0.12 vs. 0.15). Thus, comparisons limited to a single distance measure may underestimate the performance of conventional approaches. MinHash performed substantially worse than the other methods. With 128 hash values, the standard error of the estimated Jaccard similarity can reach approximately 0.044, suggesting that sketch size may contribute to its limited discrimination between fragments.

Forward fragments and their reverse complements showed similar results across methods, including conventional k-mer profiles. However, this agreement does not establish that gLMs produce strand-invariant representations. Such a conclusion would require directly comparing the representations of individual fragments with those of their own reverse complements.

A further question was whether the held-out fragments collapse into one region of the embedding space. Egoviruses and mriyaviruses are seen by the models only at inference, and sequences far from the training data can give unstable representations that look alike whatever their origin. Such a representation could still score well on genome-level proximity, since fragments of one genome would lie close together, while hiding the difference between the two orders. We therefore measured inter-order overlap, the share of mriyavirus fragments among the nearest neighbors of egovirus fragments, with one reference subsample and the mriyavirus fragments in the search space. Lower values mean better separation, and the value expected by chance equals the share of mriyavirus fragments in the search space. Read together with P@10, the metric shows whether a layer keeps the fragments of a genome together and the two orders apart. The results suggest that the models do not place all held-out fragments in a single cluster. Still, the 4-mer profile had the lowest overlap of all points (0.12), and Evo2-7B at layer 26, the layer chosen on the reference set, separated the orders worse than the baseline (0.26). The metric has its limits: it compares only two orders and uses egoviruses as queries, so it says nothing about separation from other viruses.

### Limitations

Several limitations should be considered. We evaluated only 2,000-nucleotide fragments, and performance may differ for longer contigs. The reference benchmark was constructed from curated ICTV VMR genomes and may not fully reflect the complexity of real metagenomic assemblies. In the reference dataset, nearest neighbors could include other fragments of the same genome, so P@5 partly reflected intra-genome similarity. For the held-out viruses, genome identity rather than taxonomic class was used as the label, making the two evaluations not directly comparable. We tested Evo2-7B rather than the larger 40B model and did not include several other available gLMs, such as Nucleotide Transformer [21], HyenaDNA [22], and Caduceus [23]. MinHash was evaluated only for k-mer lengths from 6 to 10 and a single sketch size.

The 50 reference subsamples were drawn from the same pool of genomes, and the held-out genomes remained unchanged across evaluations. Consequently, the statistical comparisons reflect robustness to reference subsampling rather than generalizability to other viral genomes.

We obtained the results for the four gLMs and the 4-mer profile with the following settings. The same random state was used to select genomes in every reference subsample, and each genome was cut into consecutive 2,000-nucleotide fragments starting from a random position. Held-out genomes were split into their first ten consecutive fragments and a terminal remainder. ARI was computed with the KMedoids implementation from scikit-learn-extra. For ViraLM, the two 1,000-nucleotide windows of each fragment entered the search space as separate points. For DNABERT-2 and DNABERT-S, mean pooling ran over all token positions, padding included.

## Conclusion

We evaluated four genomic language models on viral genome fragments and compared them with conventional k-mer methods and MinHash. Three of the four gLMs outperformed the k-mer baseline on the reference dataset, although the improvements were moderate. The best representations generally came from intermediate layers. On held-out egoviruses and mriyaviruses, DNABERT-S showed the most consistently strong performance, while k-mer profiles with Jensen–Shannon distance remained competitive. MinHash performed substantially worse.

These results suggest that gLMs can provide more taxonomically informative representations than conventional methods, but larger models are not necessarily better. The performance of DNABERT-S points to the potential benefits of contrastive fine-tuning.

## Supplementary Materials

The following supporting information can be downloaded at: [link to be added]. The file contains 34 supplementary figures, one per page. Figure S1: Choice of the k-mer length on the reference set: (a) ARI and precision@5 of the k-mer profile with the cosine distance as a function of k; (b) precision@5 of the k-mer profile with the cosine and Jensen–Shannon distances and of MinHash sketches (mean ± standard deviation over the reference subsets). Figure S2: Precision@5 of each method on the held-out viruses over the 50 reference subsets (boxes show the median and quartiles, points show subsets; the layer is chosen on the reference set), with the Benjamini–Hochberg-adjusted significance of paired Wilcoxon tests between neighboring methods. Figures S3–S14: Layer-wise ARI (S3–S6), precision@5 (S7–S10) and silhouette coefficient (S11–S14) of DNABERT-2, DNABERT-S, ViraLM and Evo2-7B on the reference dataset. Figures S15–S30: Layer-wise precision@5 of the same four models on egoviruses and mriyaviruses and on their reverse complements. Figures S31–S34: Statistical comparison of the methods over the reference subsamples: score distributions and paired differences for the reference benchmark (S31, S32), paired differences for the held-out viruses (S33) and for reverse-complement proximity (S34). Table S1: Evo2-7B at layer 3 and at layer 26 on the held-out viruses.

## Data and Code Availability

The benchmark is based on ICTV VMR (https://ictv.global/vmr). The processing pipeline, metric code, and notebook for generating images can be found at https://github.com/tse-kladman/glm_vectorizing. The pre-trained models are publicly available: DNABERT-2 (https://huggingface.co/zhihan1996/DNABERT-2-117M), DNABERT-S (https://huggingface.co/zhihan1996/DNABERT-S), ViraLM (https://github.com/ChengPENG-wolf/ViraLM), and Evo2 (https://github.com/ArcInstitute/evo2). The layer and k-mer length chosen for each method are stored in results/metrics/selection.json, and the metric values for every reference subset are included in the repository.

## References

1. Peng, C.; Shang, J.; Guan, J.; Wang, D.; Sun, Y. ViraLM: Empowering Virus Discovery through the Genome Foundation Model. Bioinformatics 2024, 40, doi:10.1093/bioinformatics/btae704.

2. Benegas, G.; Ye, C.; Albors, C.; Li, J.C.; Song, Y.S. Genomic Language Models: Opportunities and Challenges. Trends Genet. 2025, 41, 286–302, doi:10.1016/j.tig.2024.11.013.

3. Hou, X.; He, Y.; Fang, P.; Mei, S.-Q.; Xu, Z.; Wu, W.-C.; Tian, J.-H.; Zhang, S.; Zeng, Z.-Y.; Gou, Q.-Y.; et al. Using Artificial Intelligence to Document the Hidden RNA Virosphere. Cell 2024, 187, 6929-6942.e16, doi:10.1016/j.cell.2024.09.027.

4. International Committee on Taxonomy of Viruses (ICTV) Virus Metadata Resource (VMR), MSL40 V2 2025.

5. Gaïa, M.; Guyet, U.; Ruscheweyh, H.-J.; Eren, A.M.; Sunagawa, S.; Koonin, E.V.; Krupovic, M.; Delmont, T.O. Closest Relatives of Poxviruses Replicate in the Digestive System of Humans and Animals Worldwide. 2024, doi:10.1101/2024.03.23.586382.

6. Yutin, N.; Mutz, P.; Krupovic, M.; Koonin, E.V. Mriyaviruses: Small Relatives of Giant Viruses. mBio 2024, 15, doi:10.1128/mbio.01035-24.

7. Zhou, Z.; Ji, Y.; Li, W.; Dutta, P.; Davuluri, R.; Liu, H. DNABERT-2: Efficient Foundation Model and Benchmark For Multi-Species Genome 2023.

8. Zhou, Z.; Wu, W.; Ho, H.; Wang, J.; Shi, L.; Davuluri, R.V.; Wang, Z.; Liu, H. DNABERT-S: Pioneering Species Differentiation with Species-Aware DNA Embeddings. Bioinformatics 2025, 41, i255–i264, doi:10.1093/bioinformatics/btaf188.

9. Brixi, G.; Durrant, M.G.; Ku, J.; Naghipourfar, M.; Poli, M.; Sun, G.; Brockman, G.; Chang, D.; Fanton, A.; Gonzalez, G.A.; et al. Genome Modelling and Design across All Domains of Life with Evo 2. Nature 2026, 652, 1349–1361, doi:10.1038/s41586-026-10176-5.

10. Lin, J. Divergence Measures Based on the Shannon Entropy. IEEE Trans. Inf. Theory 1991, 37, 145–151, doi:10.1109/18.61115.

11. Endres, D.M.; Schindelin, J.E. A New Metric for Probability Distributions. IEEE Trans. Inf. Theory 2003, 49, 1858–1860, doi:10.1109/tit.2003.813506.

12. Broder, A.Z. On the Resemblance and Containment of Documents. In Proceedings of the Proceedings. Compression and Complexity of SEQUENCES 1997 (Cat. No.97TB100171); IEEE Comput. Soc; pp. 21–29.

13. Ondov, B.D.; Treangen, T.J.; Melsted, P.; Mallonee, A.B.; Bergman, N.H.; Koren, S.; Phillippy, A.M. Mash: Fast Genome and Metagenome Distance Estimation Using MinHash. Genome Biol. 2016, 17, doi:10.1186/s13059-016-0997-x.

14. Ji, Y.; Zhou, Z.; Liu, H.; Davuluri, R.V. DNABERT: Pre-Trained Bidirectional Encoder Representations from Transformers Model for DNA-Language in Genome. Bioinformatics 2021, 37, 2112–2120, doi:10.1093/bioinformatics/btab083.

15. Vaswani, A.; Shazeer, N.; Parmar, N.; Uszkoreit, J.; Jones, L.; Gomez, A.N.; Kaiser, L.; Polosukhin, I. Attention Is All You Need 2017.

16. Hubert, L.; Arabie, P. Comparing Partitions. J. Classif. 1985, 2, 193–218, doi:10.1007/bf01908075.

17. Kaufman, L.; Rousseeuw, P.J. Finding Groups in Data: An Introduction to Cluster Analysis; Wiley, 1990; ISBN 978-0-470-31680-1.

18. Rousseeuw, P.J. Silhouettes: A Graphical Aid to the Interpretation and Validation of Cluster Analysis. J. Comput. Appl. Math. 1987, 20, 53–65, doi:10.1016/0377-0427(87)90125-7.

19. Wilcoxon, F. Individual Comparisons by Ranking Methods. Biom. Bull. 1945, 1, 80, doi:10.2307/3001968.

20. Benjamini, Y.; Hochberg, Y. Controlling the False Discovery Rate: A Practical and Powerful Approach to Multiple Testing. J. R. Stat. Soc. Ser. B Stat. Methodol. 1995, 57, 289–300, doi:10.1111/j.2517-6161.1995.tb02031.x.

21. Dalla-Torre, H.; Gonzalez, L.; Mendoza-Revilla, J.; Lopez Carranza, N.; Grzywaczewski, A.H.; Oteri, F.; Dallago, C.; Trop, E.; de Almeida, B.P.; Sirelkhatim, H.; et al. Nucleotide Transformer: Building and Evaluating Robust Foundation Models for Human Genomics. Nat. Methods 2024, 22, 287–297, doi:10.1038/s41592-024-02523-z.

22. Nguyen, E.; Poli, M.; Faizi, M.; Thomas, A.; Birch-Sykes, C.; Wornow, M.; Patel, A.; Rabideau, C.; Massaroli, S.; Bengio, Y.; et al. HyenaDNA: Long-Range Genomic Sequence Modeling at Single Nucleotide Resolution 2023.

23. Schiff, Y.; Kao, C.-H.; Gokaslan, A.; Dao, T.; Gu, A.; Kuleshov, V. Caduceus: Bi-Directional Equivariant Long-Range DNA Sequence Modeling 2024.

