## Supplementary figures for "Assessing the Capacity of Genomic Language Models to Generate Informative Embeddings of Viral Genome Fragments"

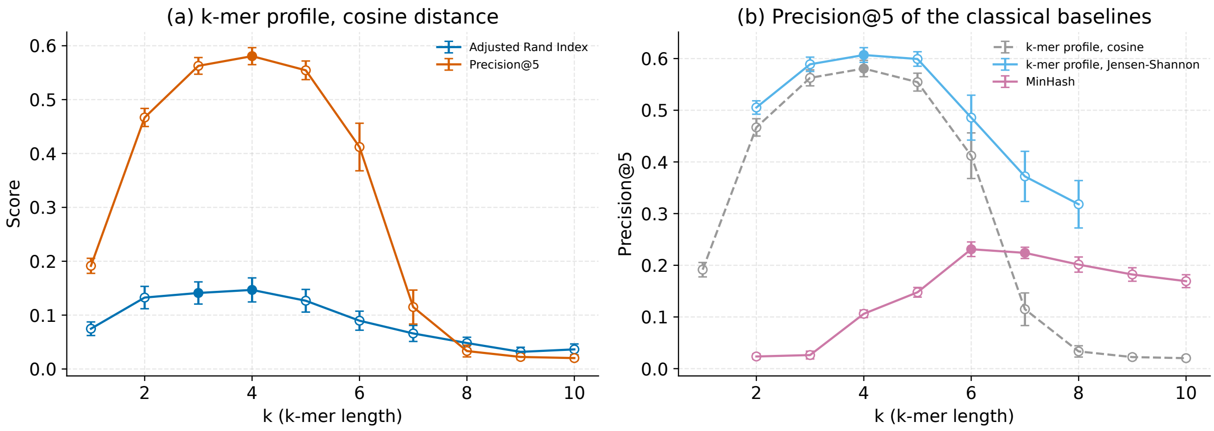


**Supplementary Figure S1.** Choice of the k-mer length on the reference set. (a) ARI and precision@5 of the k-mer profile with the cosine distance as a function of the k-mer length k. (b) Precision@5 of the k-mer profile with the cosine and with the Jensen–Shannon distance and of MinHash sketches. Points show the mean ± standard deviation over the reference subsets (50 subsets for the cosine distance, k = 1–10; 10 subsets for the Jensen–Shannon distance, k = 2–8, and for MinHash, k = 2–10). Filled markers indicate the best k of each curve and the values that do not differ significantly from it; open markers indicate k values significantly worse than the best k of the same curve (Benjamini–Hochberg-adjusted $p < 0.05$, paired Wilcoxon test).

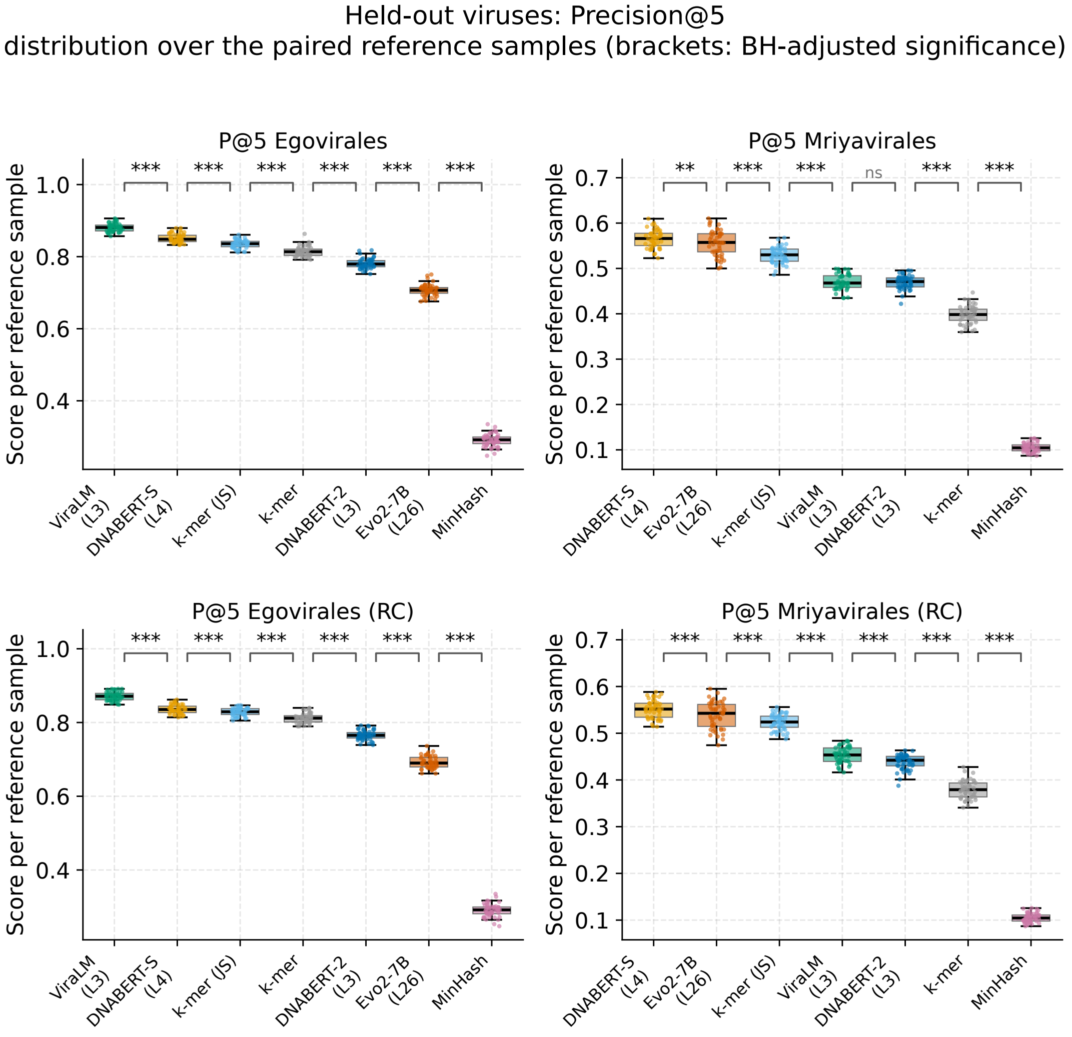


**Supplementary Figure S2.** Precision@5 of each method on the held-out viruses over the 50 reference subsets, for egoviruses and mriyaviruses and their reverse complements (RC). Boxes show the median and quartiles, points show individual subsets; the layer of each model (L) is chosen on the reference set. Methods are ordered by their mean score. Brackets between neighboring methods show Benjamini–Hochberg-adjusted p-values of two-sided paired Wilcoxon tests: * $p < 0.05$, ** $p < 0.01$, *** $p < 0.001$; ns, not significant.

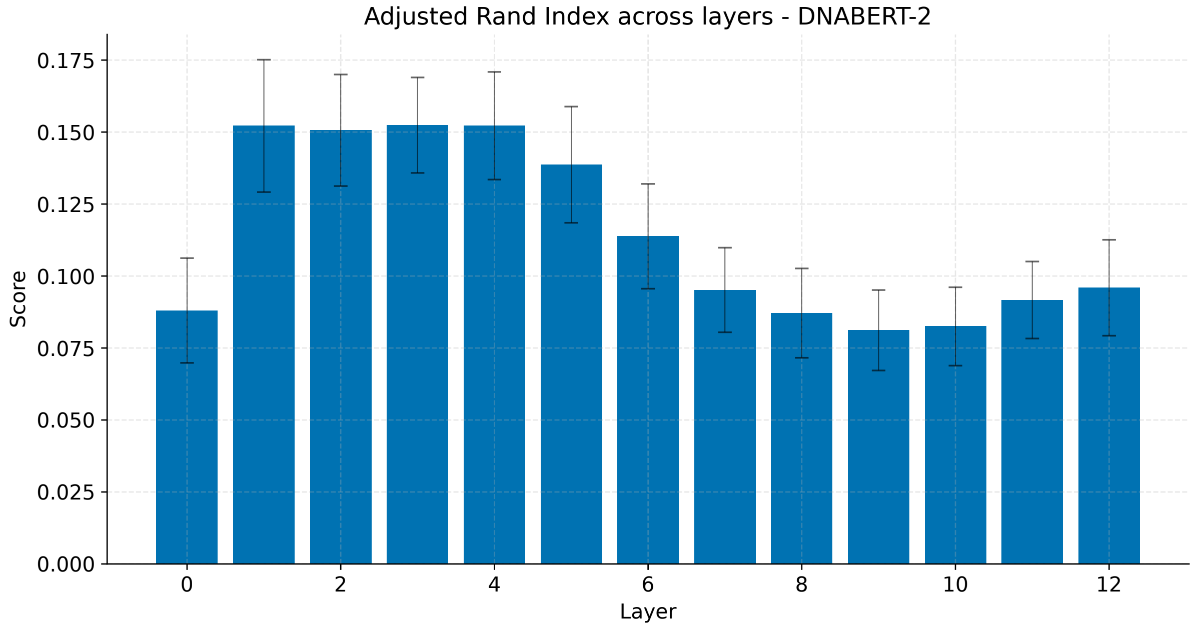


**Supplementary Figure S3.** Adjusted Rand index (ARI) on the reference dataset for each layer of DNABERT-2. Layers are numbered from 0 (embedding layer) to 12 (last transformer block). Bars show the mean ± standard deviation across the 50 reference subsamples.

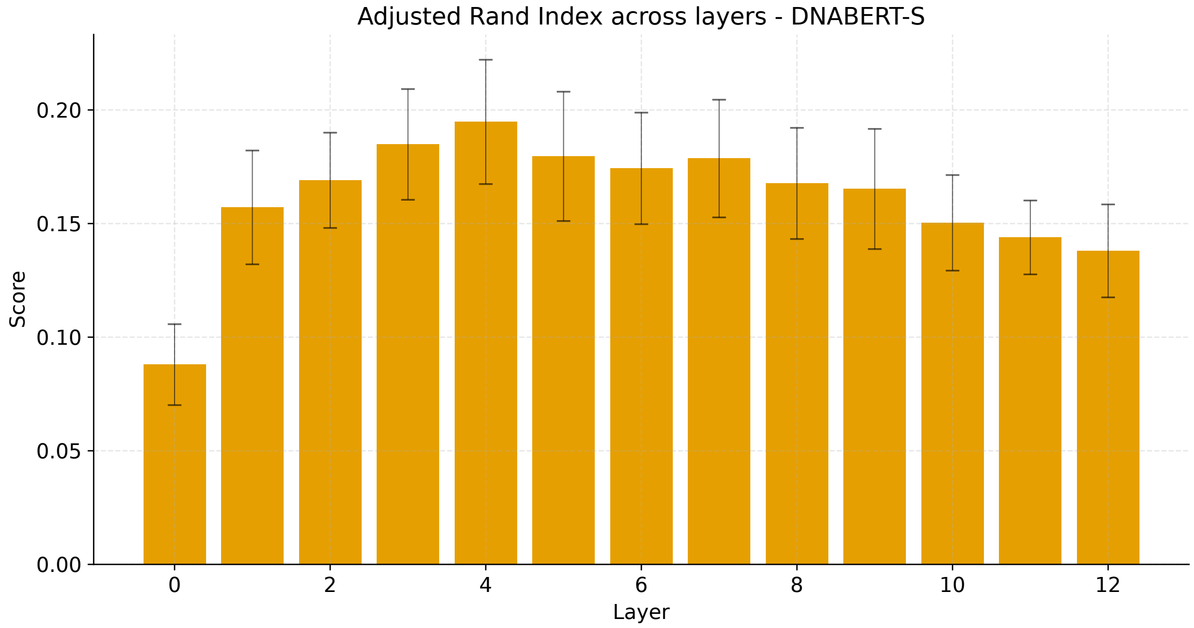


**Supplementary Figure S4.** Adjusted Rand index (ARI) on the reference dataset for each layer of DNABERT-S. Layers are numbered from 0 (embedding layer) to 12 (last transformer block). Bars show the mean ± standard deviation across the 50 reference subsamples.

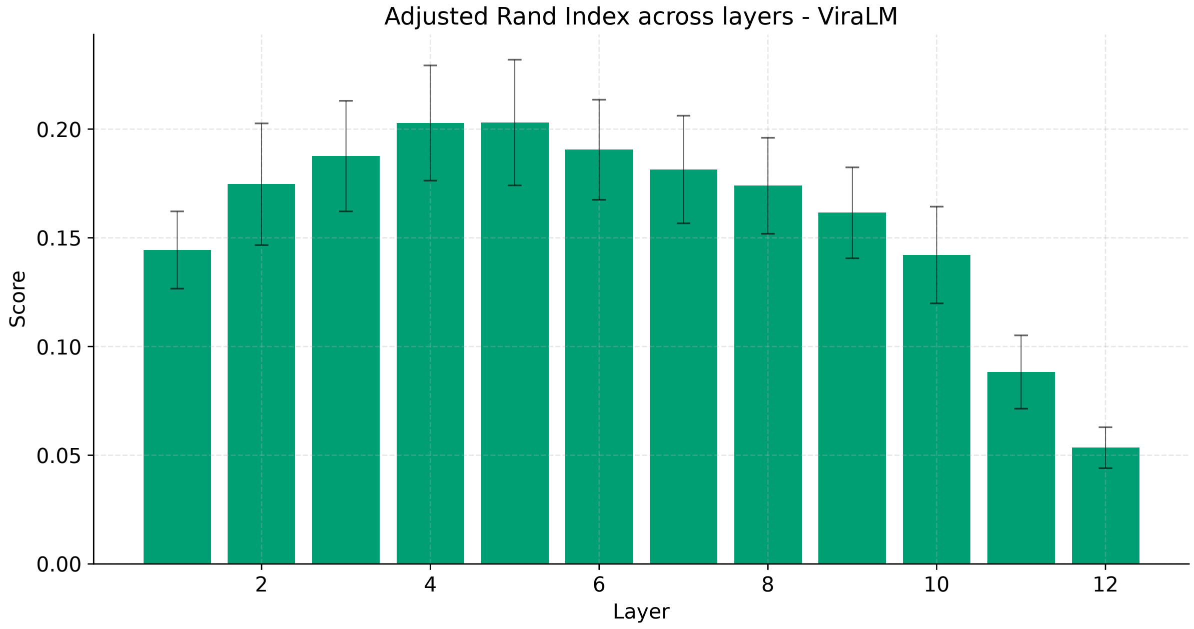


**Supplementary Figure S5.** Adjusted Rand index (ARI) on the reference dataset for each layer of ViraLM. Layers are the transformer blocks 1–12. Bars show the mean ± standard deviation across the 50 reference subsamples.

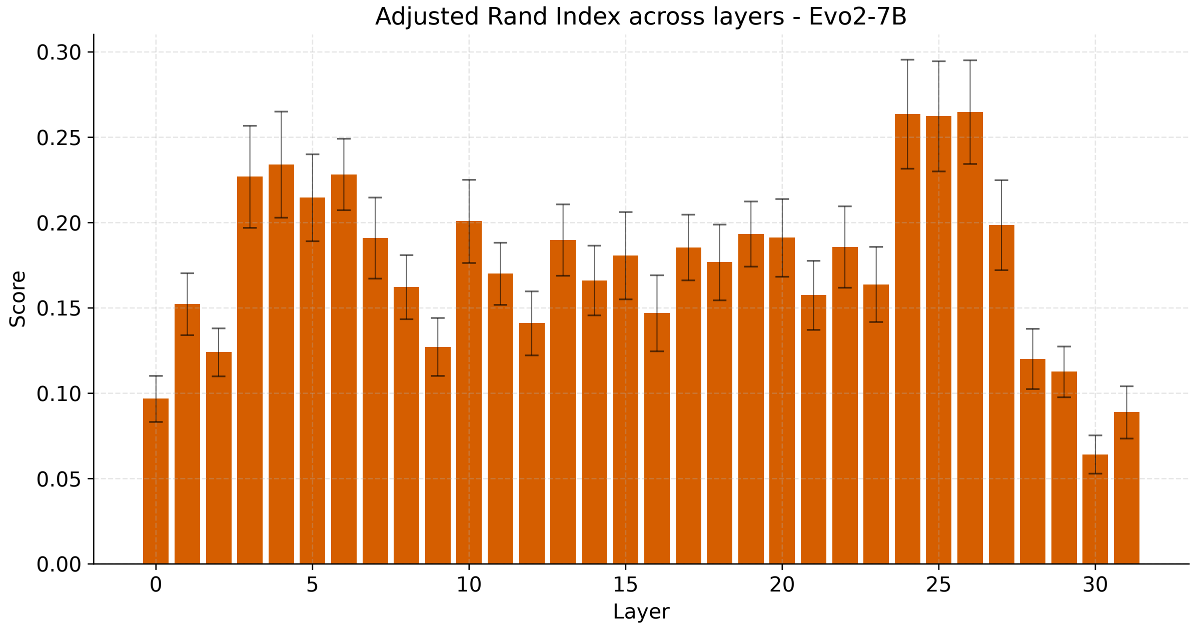


**Supplementary Figure S6.** Adjusted Rand index (ARI) on the reference dataset for each layer of Evo2-7B. Layers are the 32 blocks of the model, numbered 0–31. Bars show the mean ± standard deviation across the 50 reference subsamples.

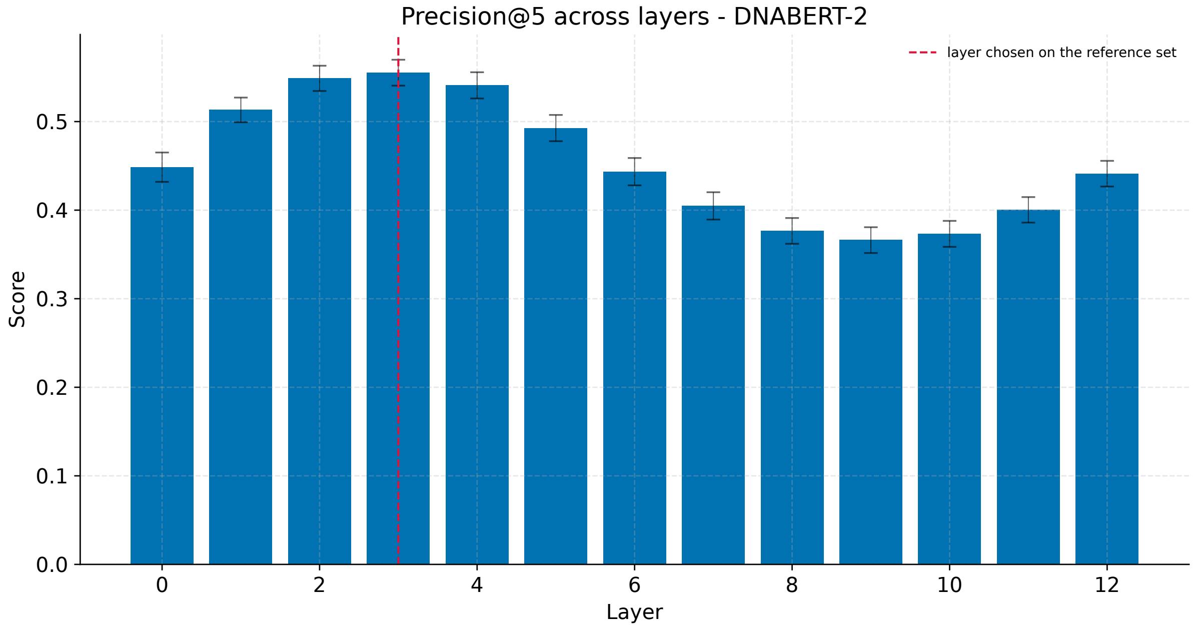


**Supplementary Figure S7.** Precision@5 (P@5) on the reference dataset for each layer of DNABERT-2. Layers are numbered from 0 (embedding layer) to 12 (last transformer block). Bars show the mean ± standard deviation across the 50 reference subsamples. The dashed line marks the layer chosen on the reference set.

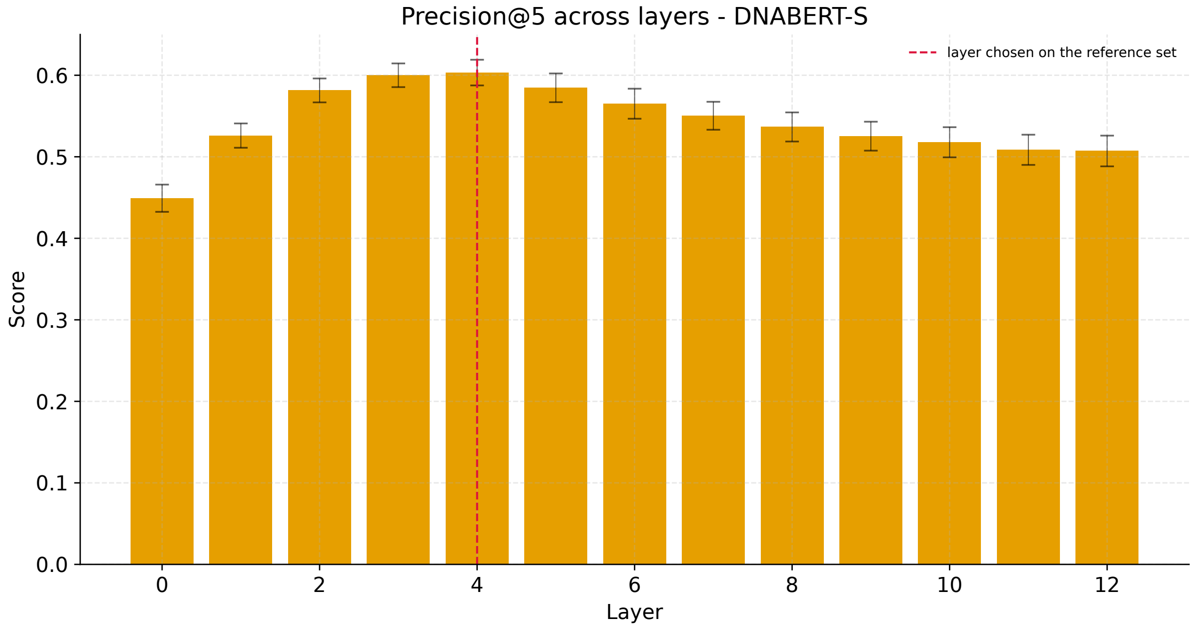


**Supplementary Figure S8.** Precision@5 (P@5) on the reference dataset for each layer of DNABERT-S. Layers are numbered from 0 (embedding layer) to 12 (last transformer block). Bars show the mean ± standard deviation across the 50 reference subsamples. The dashed line marks the layer chosen on the reference set.

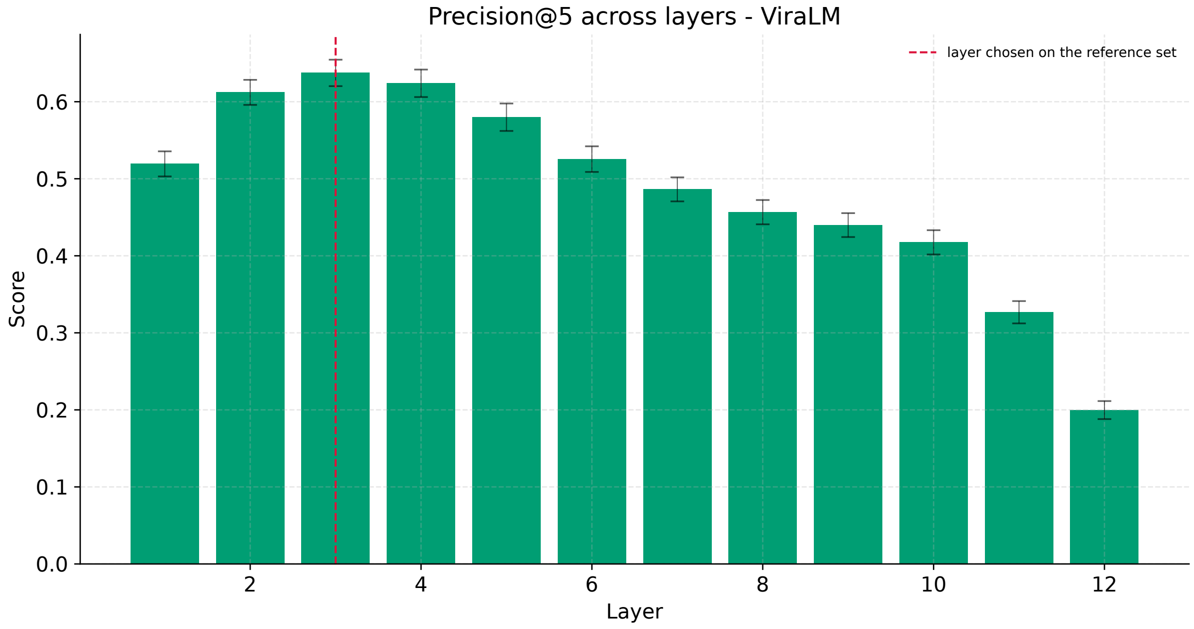


**Supplementary Figure S9.** Precision@5 (P@5) on the reference dataset for each layer of ViraLM. Layers are the transformer blocks 1–12. Bars show the mean ± standard deviation across the 50 reference subsamples. The dashed line marks the layer chosen on the reference set.

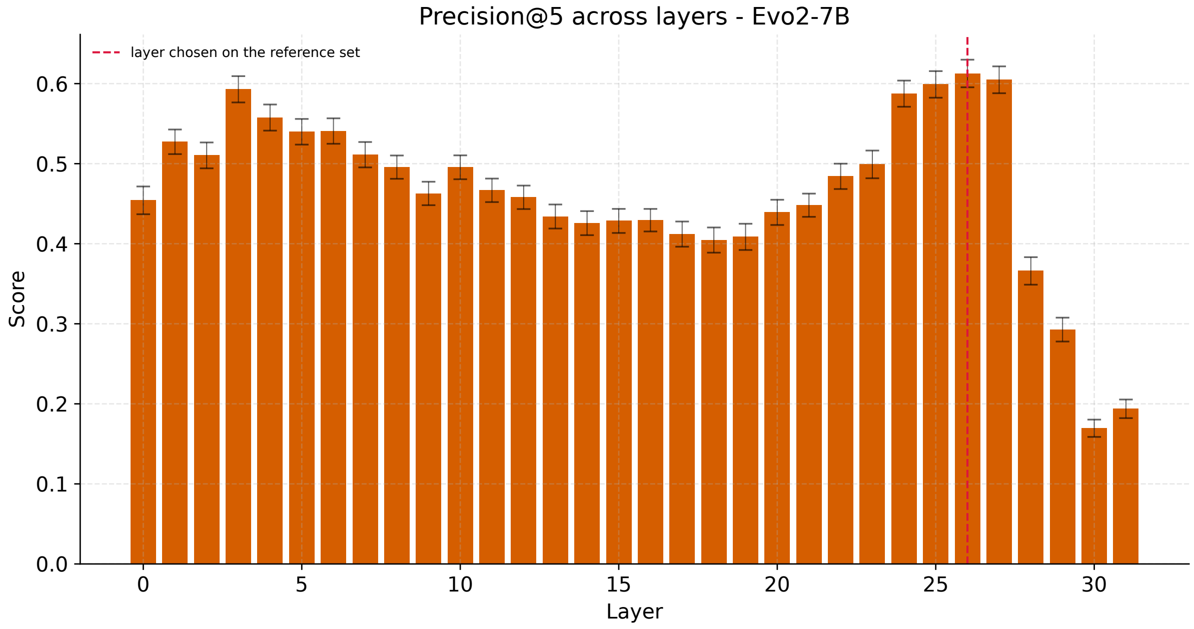


**Supplementary Figure S10.** Precision@5 (P@5) on the reference dataset for each layer of Evo2-7B. Layers are the 32 blocks of the model, numbered 0–31. Bars show the mean ± standard deviation across the 50 reference subsamples. The dashed line marks the layer chosen on the reference set.

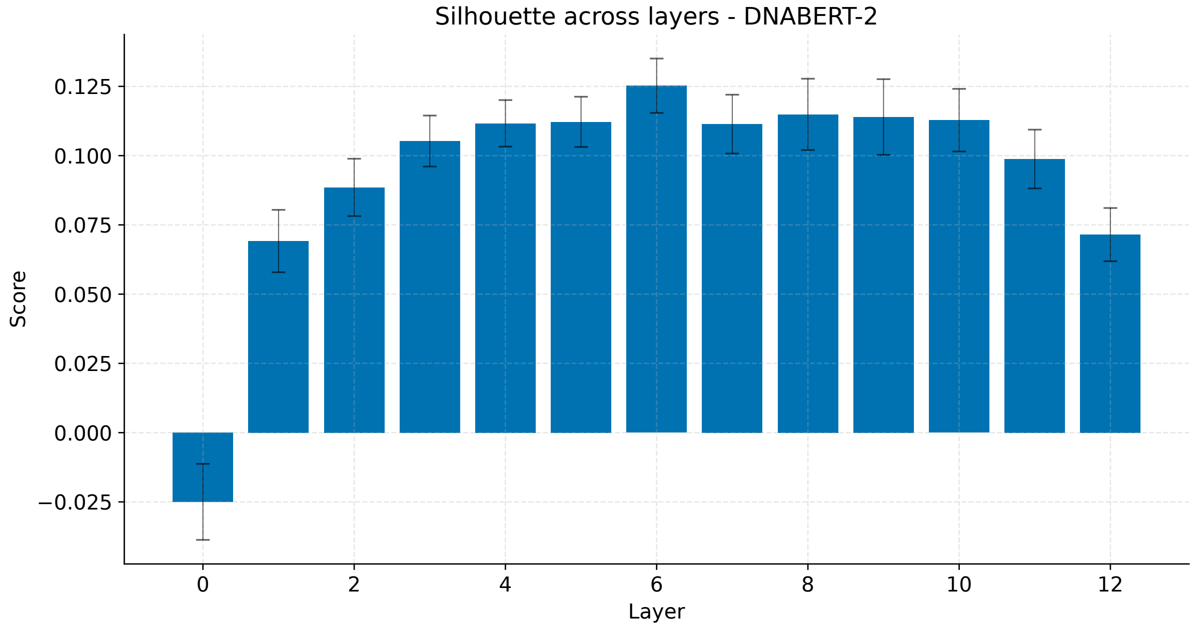


**Supplementary Figure S11.** Silhouette coefficient on the reference dataset for each layer of DNABERT-2. Layers are numbered from 0 (embedding layer) to 12 (last transformer block). Bars show the mean ± standard deviation across the 50 reference subsamples.

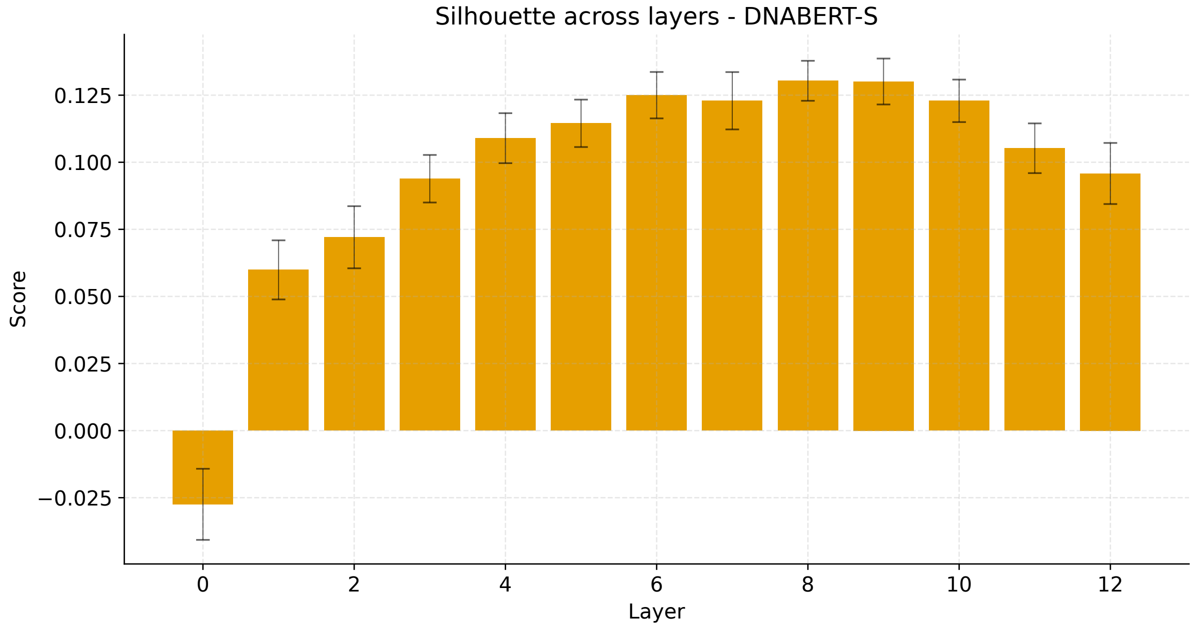


**Supplementary Figure S12.** Silhouette coefficient on the reference dataset for each layer of DNABERT-S. Layers are numbered from 0 (embedding layer) to 12 (last transformer block). Bars show the mean ± standard deviation across the 50 reference subsamples.

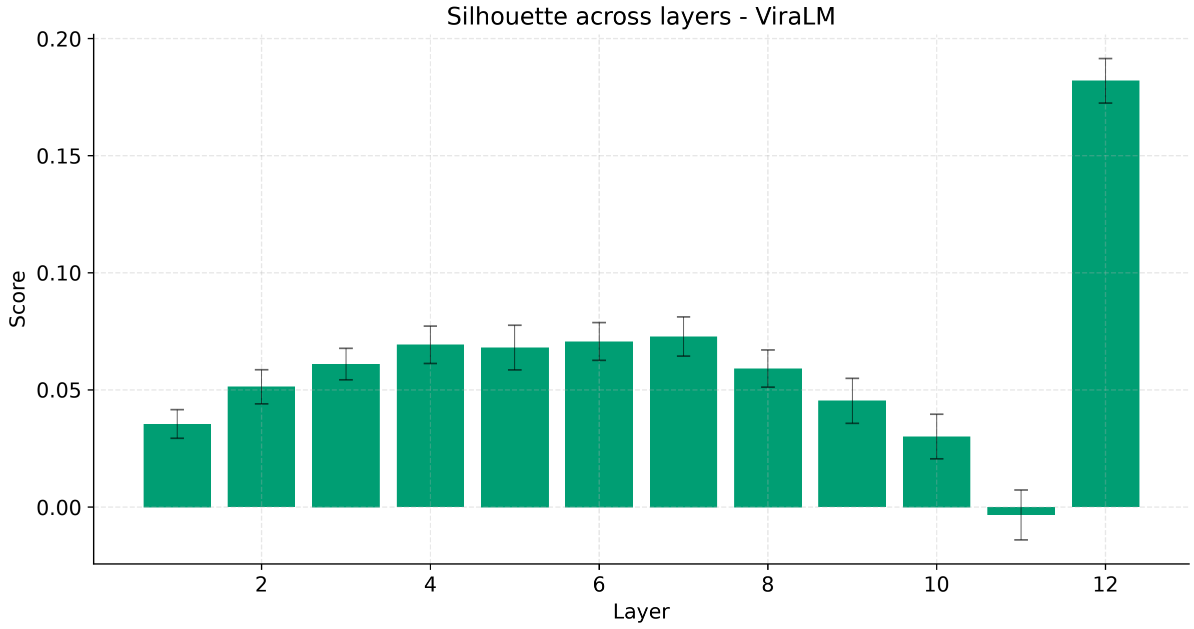


**Supplementary Figure S13.** Silhouette coefficient on the reference dataset for each layer of ViraLM. Layers are the transformer blocks 1–12. Bars show the mean ± standard deviation across the 50 reference subsamples.

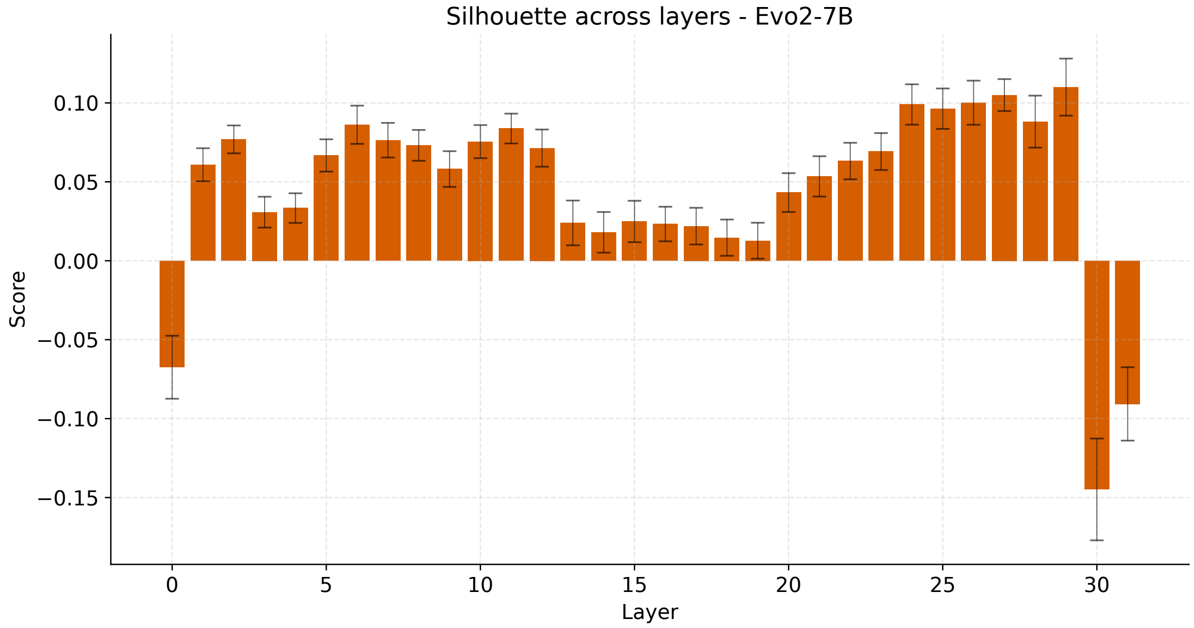


**Supplementary Figure S14.** Silhouette coefficient on the reference dataset for each layer of Evo2-7B. Layers are the 32 blocks of the model, numbered 0–31. Bars show the mean ± standard deviation across the 50 reference subsamples.

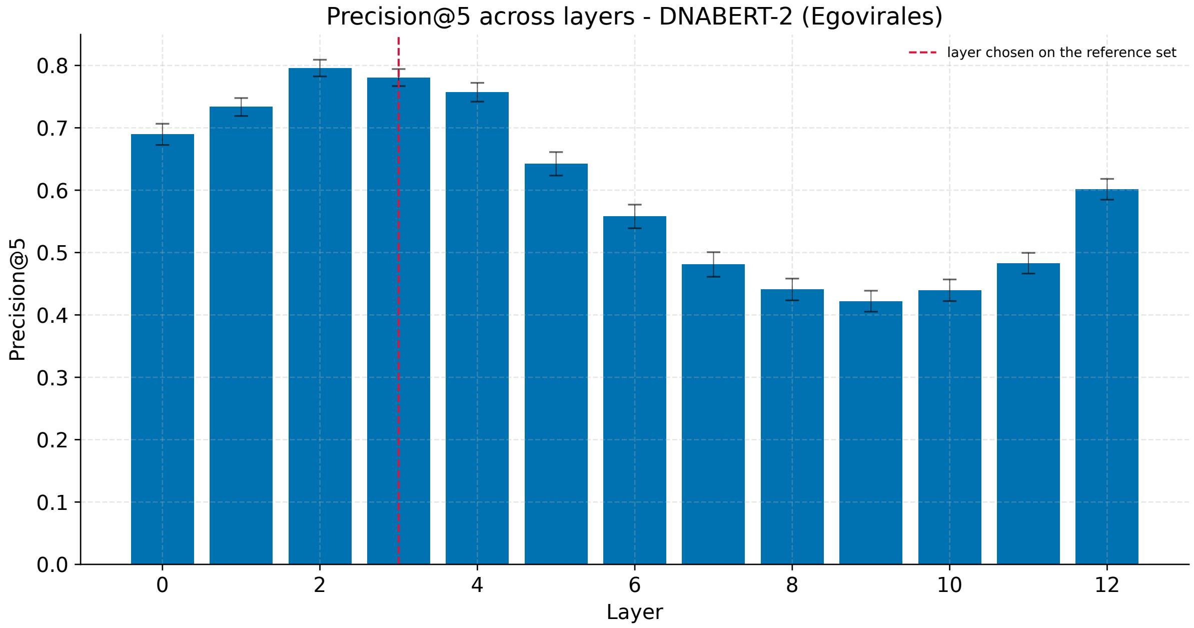


**Supplementary Figure S15.** Precision@5 on egoviruses for each layer of DNABERT-2. Precision@5 was calculated using genome identity as the label, with the search space consisting of fragments of the corresponding genome and one reference subsample. Layers are numbered from 0 (embedding layer) to 12 (last transformer block). Bars show the mean ± standard deviation across the 50 reference subsamples. The dashed line marks the layer chosen on the reference set.

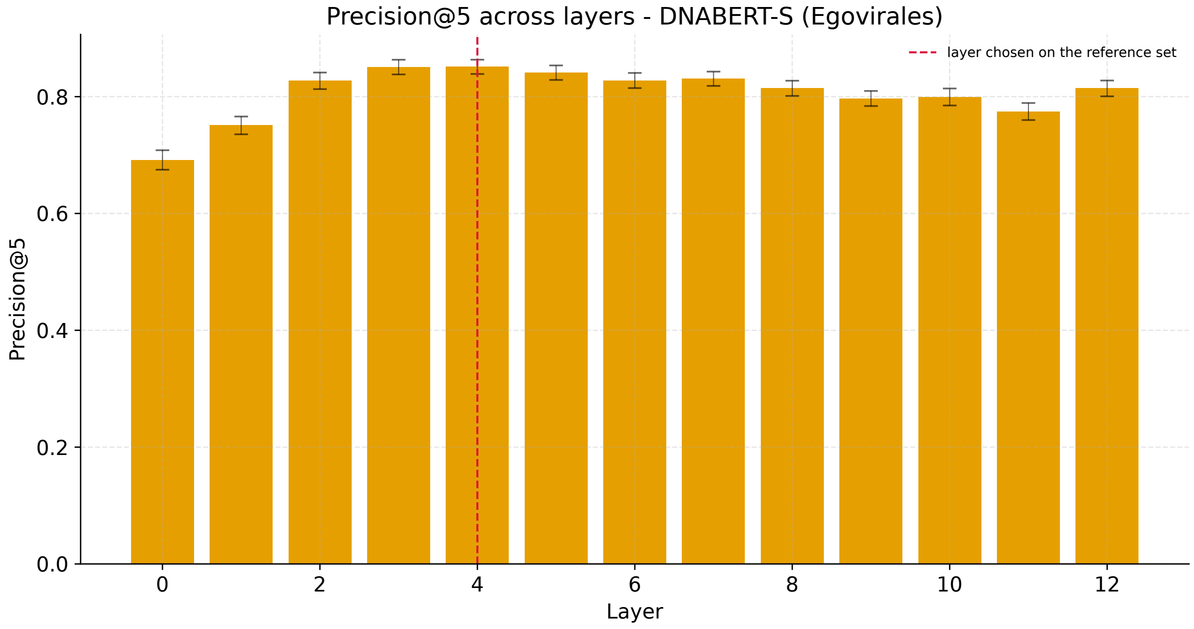


**Supplementary Figure S16.** Precision@5 on egoviruses for each layer of DNABERT-S. Precision@5 was calculated using genome identity as the label, with the search space consisting of fragments of the corresponding genome and one reference subsample. Layers are numbered from 0 (embedding layer) to 12 (last transformer block). Bars show the mean ± standard deviation across the 50 reference subsamples. The dashed line marks the layer chosen on the reference set.

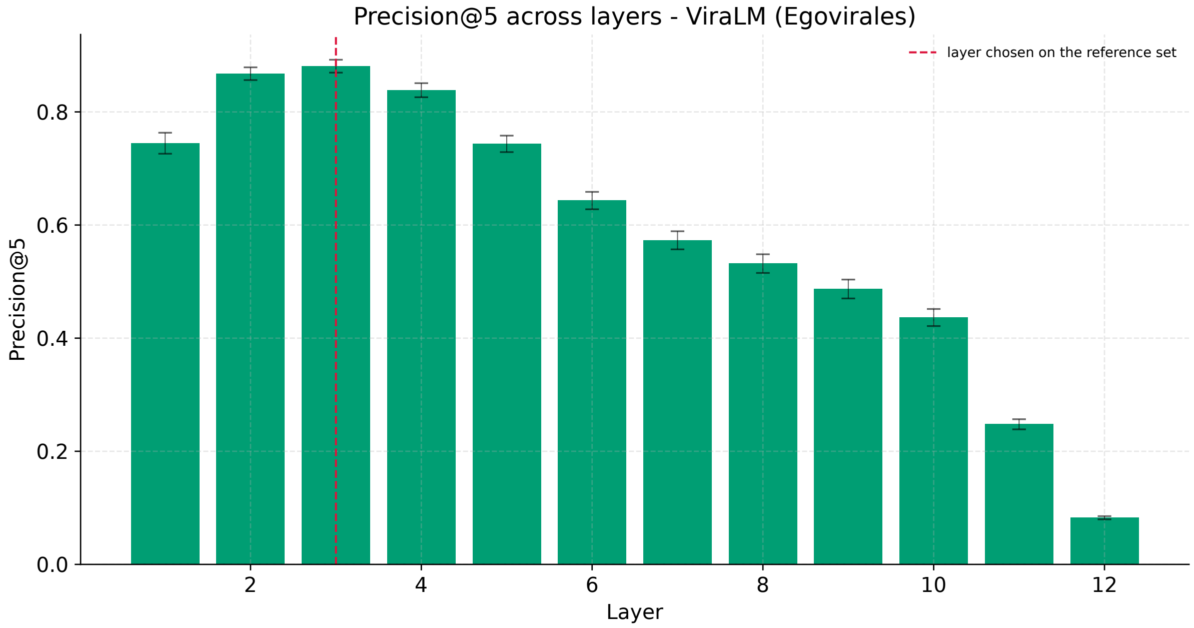


**Supplementary Figure S17.** Precision@5 on egoviruses for each layer of ViraLM. Precision@5 was calculated using genome identity as the label, with the search space consisting of fragments of the corresponding genome and one reference subsample. Layers are the transformer blocks 1–12. Bars show the mean ± standard deviation across the 50 reference subsamples. The dashed line marks the layer chosen on the reference set.

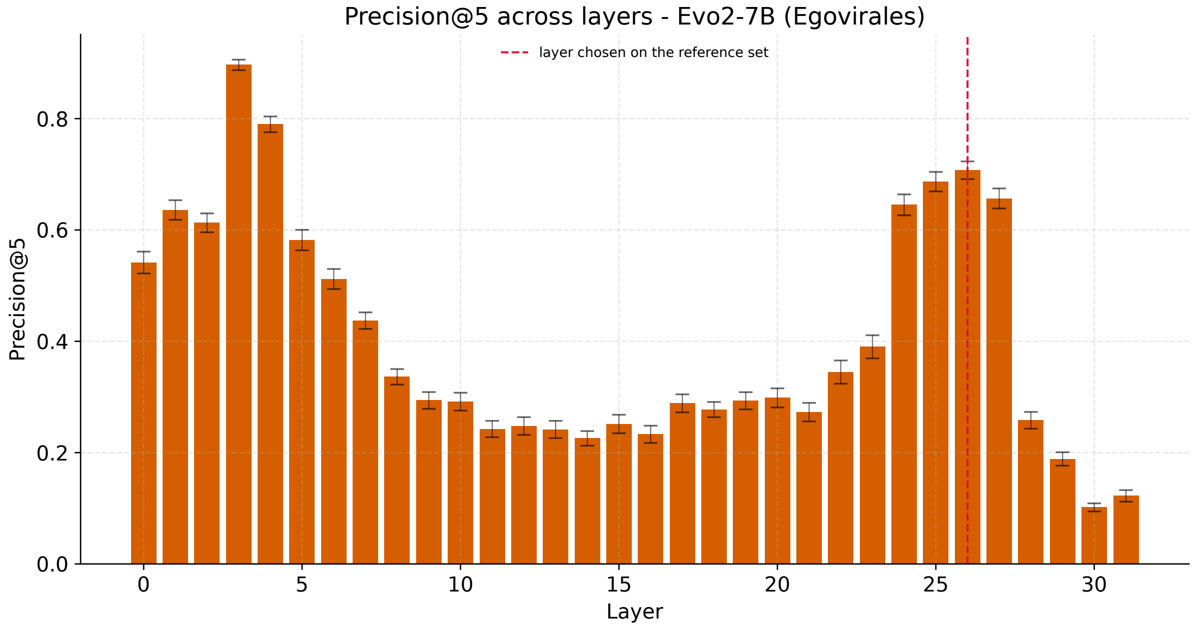


**Supplementary Figure S18.** Precision@5 on egoviruses for each layer of Evo2-7B. Precision@5 was calculated using genome identity as the label, with the search space consisting of fragments of the corresponding genome and one reference subsample. Layers are the 32 blocks of the model, numbered 0–31. Bars show the mean ± standard deviation across the 50 reference subsamples. The dashed line marks the layer chosen on the reference set.

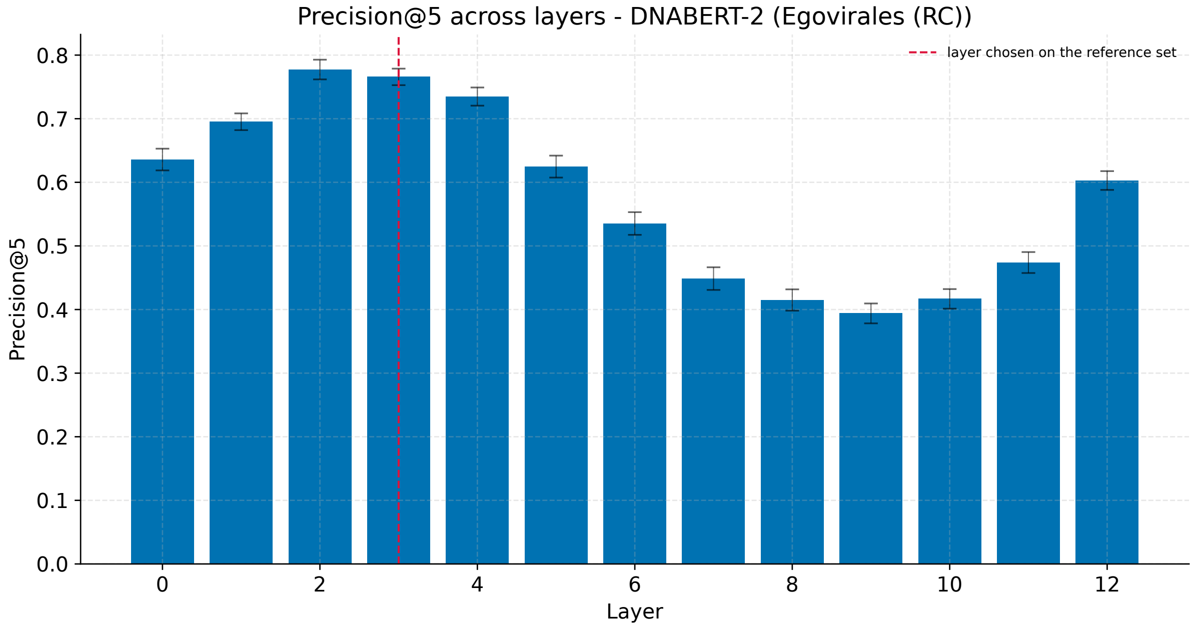


**Supplementary Figure S19.** Precision@5 on the reverse complements of egoviruses (RC) for each layer of DNABERT-2. Precision@5 was calculated using genome identity as the label, with the search space consisting of fragments of the corresponding genome and one reference subsample. Layers are numbered from 0 (embedding layer) to 12 (last transformer block). Bars show the mean ± standard deviation across the 50 reference subsamples. The dashed line marks the layer chosen on the reference set.

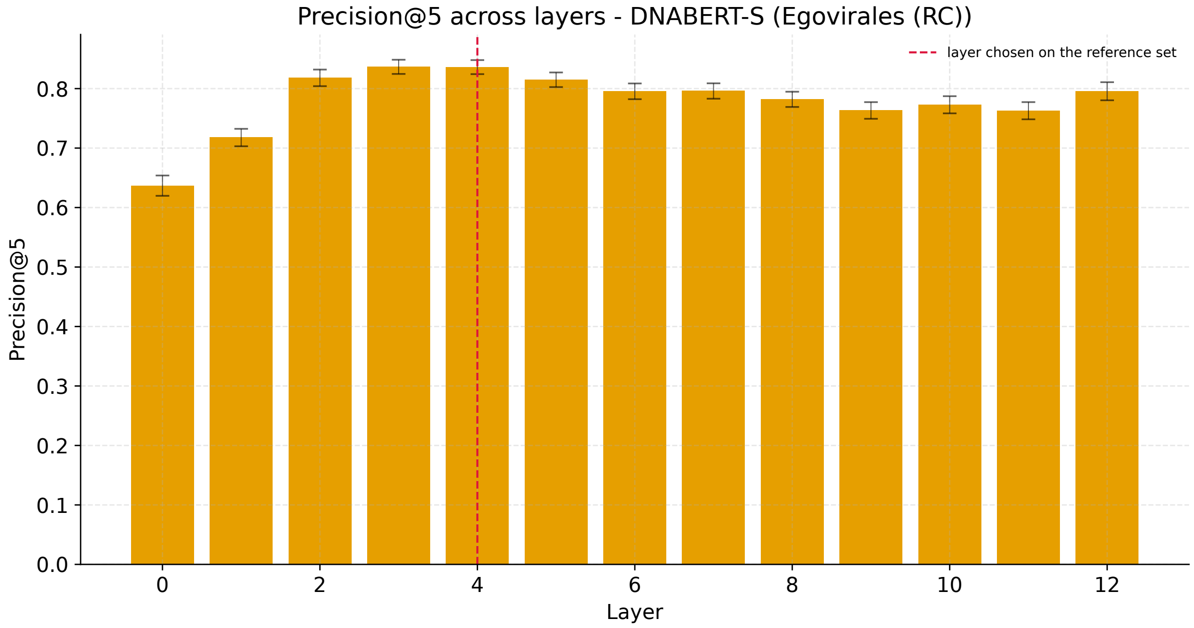


**Supplementary Figure S20.** Precision@5 on the reverse complements of egoviruses (RC) for each layer of DNABERT-S. Precision@5 was calculated using genome identity as the label, with the search space consisting of fragments of the corresponding genome and one reference subsample. Layers are numbered from 0 (embedding layer) to 12 (last transformer block). Bars show the mean ± standard deviation across the 50 reference subsamples. The dashed line marks the layer chosen on the reference set.

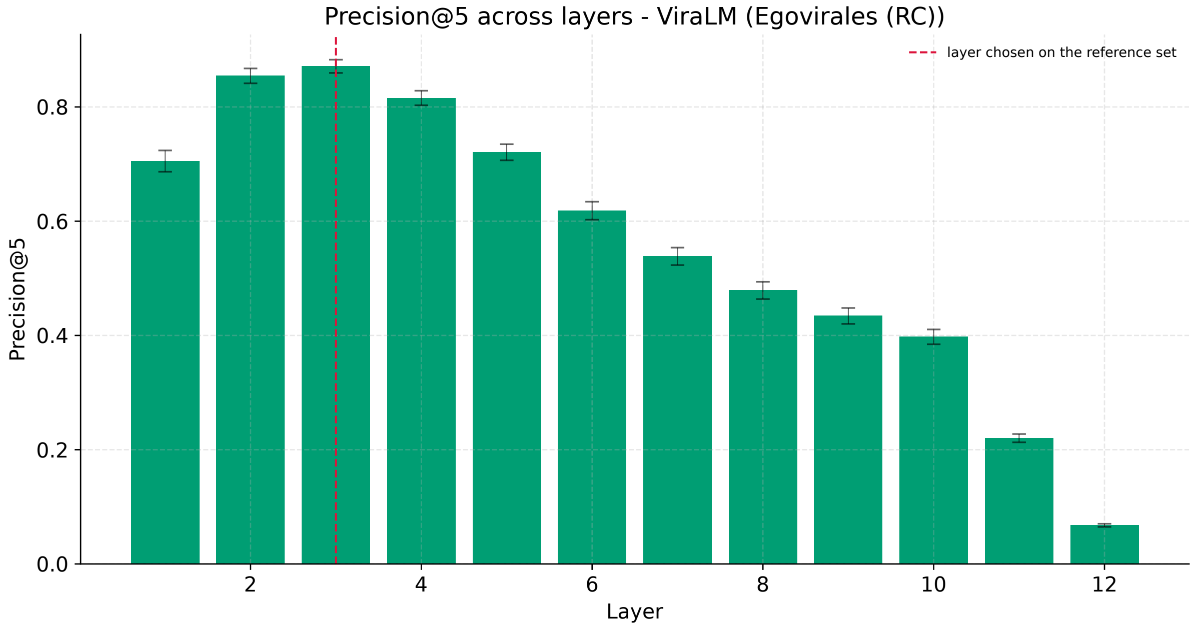


**Supplementary Figure S21.** Precision@5 on the reverse complements of egoviruses (RC) for each layer of ViraLM. Precision@5 was calculated using genome identity as the label, with the search space consisting of fragments of the corresponding genome and one reference subsample. Layers are the transformer blocks 1–12. Bars show the mean ± standard deviation across the 50 reference subsamples. The dashed line marks the layer chosen on the reference set.

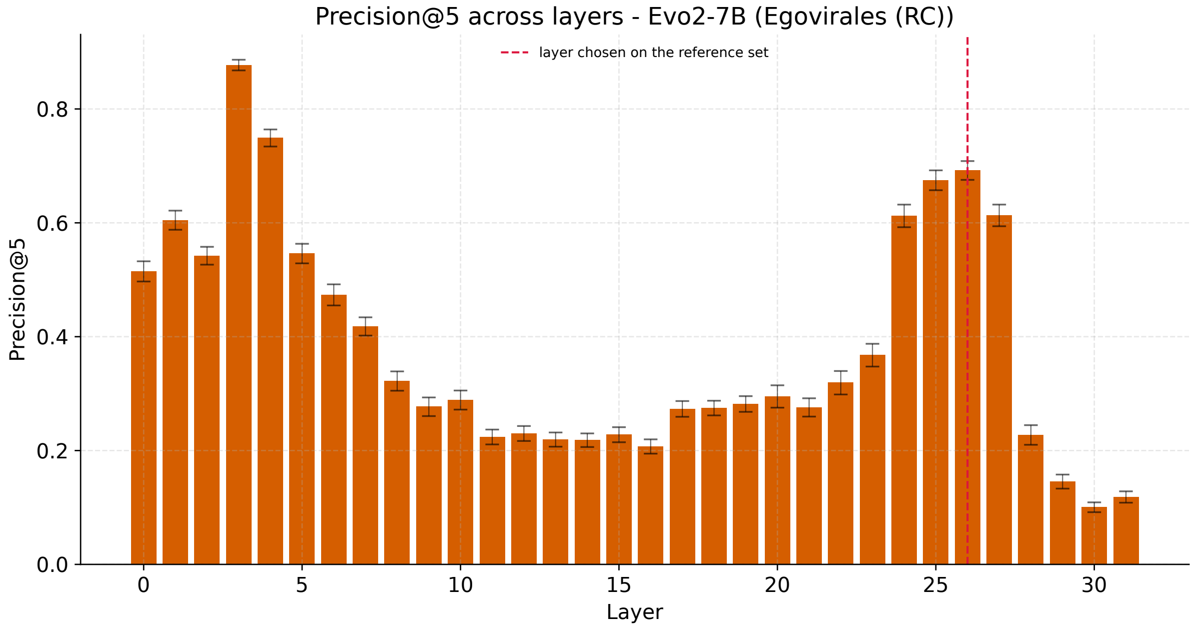


**Supplementary Figure S22.** Precision@5 on the reverse complements of egoviruses (RC) for each layer of Evo2-7B. Precision@5 was calculated using genome identity as the label, with the search space consisting of fragments of the corresponding genome and one reference subsample. Layers are the 32 blocks of the model, numbered 0–31. Bars show the mean ± standard deviation across the 50 reference subsamples. The dashed line marks the layer chosen on the reference set.

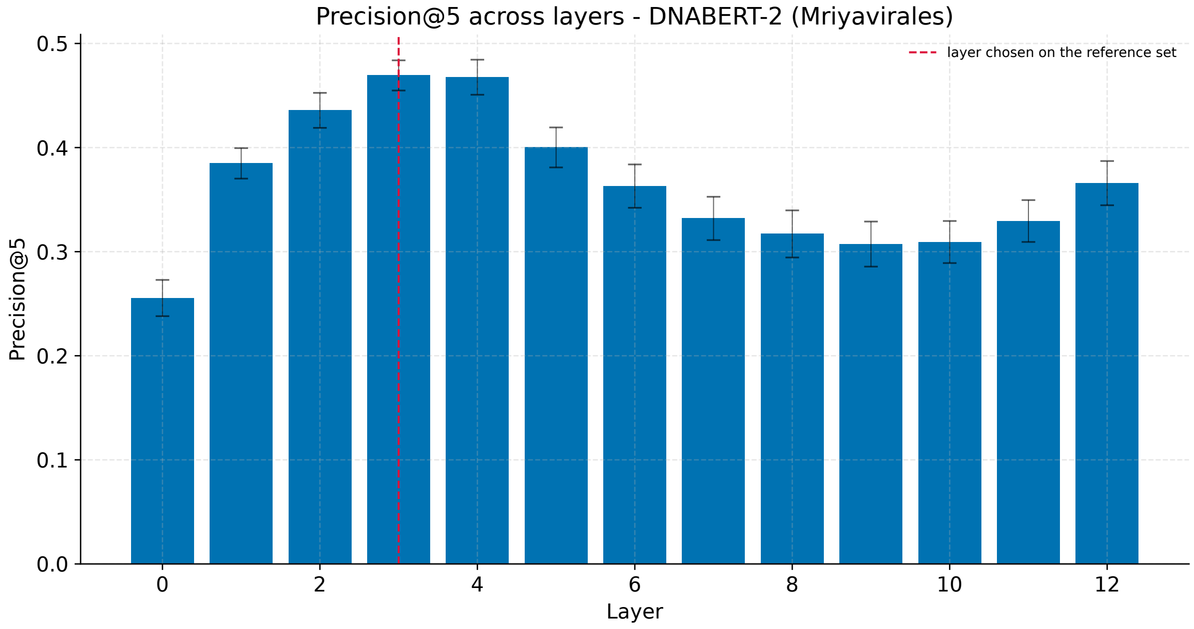


**Supplementary Figure S23.** Precision@5 on mriyaviruses for each layer of DNABERT-2. Precision@5 was calculated using genome identity as the label, with the search space consisting of fragments of the corresponding genome and one reference subsample. Layers are numbered from 0 (embedding layer) to 12 (last transformer block). Bars show the mean ± standard deviation across the 50 reference subsamples. The dashed line marks the layer chosen on the reference set.

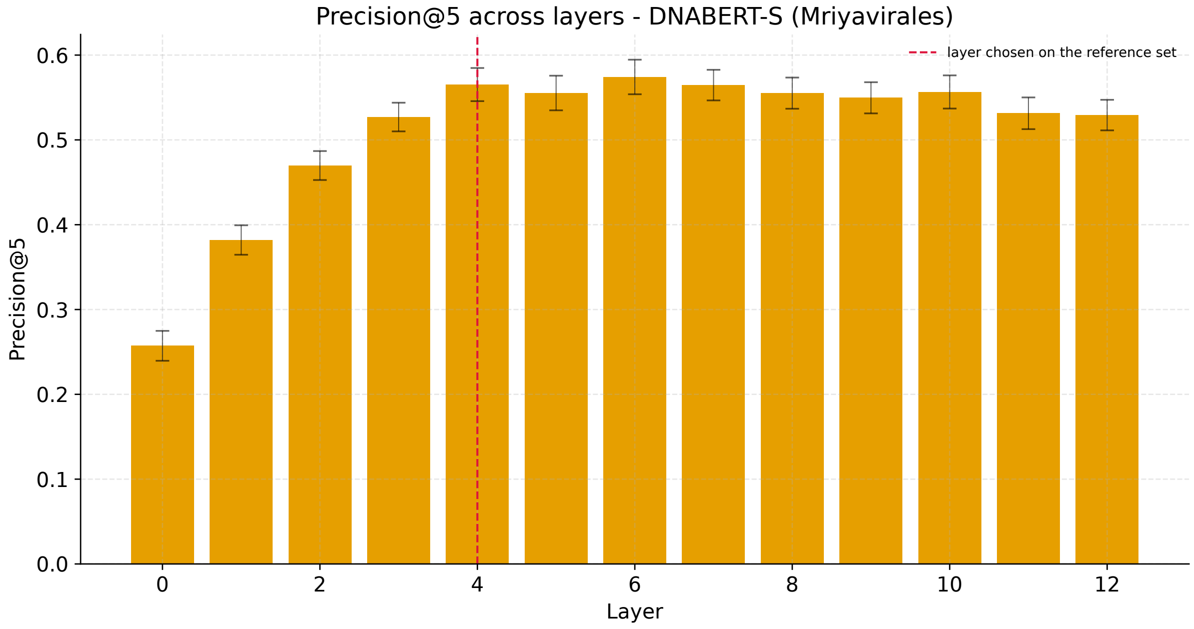


**Supplementary Figure S24.** Precision@5 on mriyaviruses for each layer of DNABERT-S. Precision@5 was calculated using genome identity as the label, with the search space consisting of fragments of the corresponding genome and one reference subsample. Layers are numbered from 0 (embedding layer) to 12 (last transformer block). Bars show the mean ± standard deviation across the 50 reference subsamples. The dashed line marks the layer chosen on the reference set.

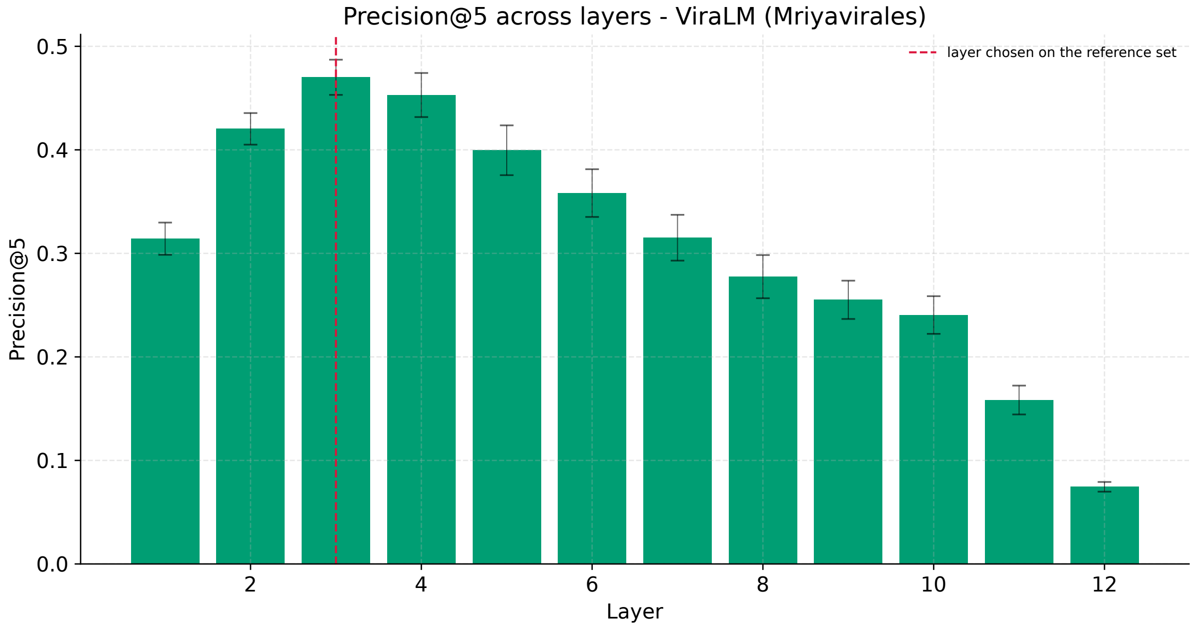


**Supplementary Figure S25.** Precision@5 on mriyaviruses for each layer of ViraLM. Precision@5 was calculated using genome identity as the label, with the search space consisting of fragments of the corresponding genome and one reference subsample. Layers are the transformer blocks 1–12. Bars show the mean ± standard deviation across the 50 reference subsamples. The dashed line marks the layer chosen on the reference set.

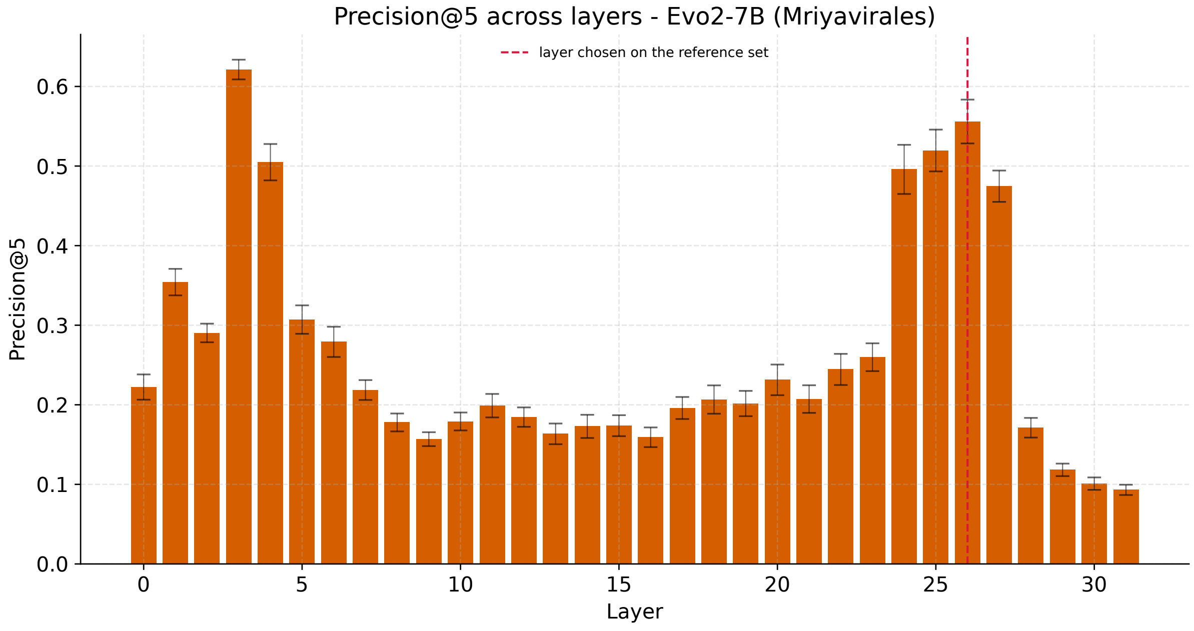


**Supplementary Figure S26.** Precision@5 on mriyaviruses for each layer of Evo2-7B. Precision@5 was calculated using genome identity as the label, with the search space consisting of fragments of the corresponding genome and one reference subsample. Layers are the 32 blocks of the model, numbered 0–31. Bars show the mean ± standard deviation across the 50 reference subsamples. The dashed line marks the layer chosen on the reference set.

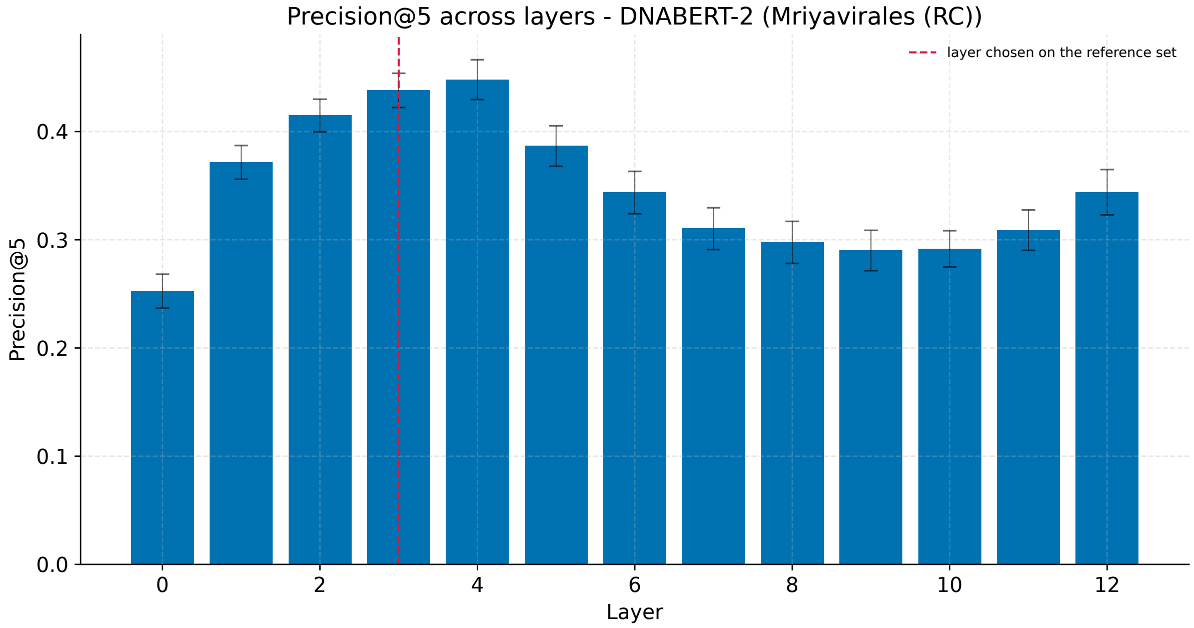


**Supplementary Figure S27.** Precision@5 on the reverse complements of mriyaviruses (RC) for each layer of DNABERT-2. Precision@5 was calculated using genome identity as the label, with the search space consisting of fragments of the corresponding genome and one reference subsample. Layers are numbered from 0 (embedding layer) to 12 (last transformer block). Bars show the mean ± standard deviation across the 50 reference subsamples. The dashed line marks the layer chosen on the reference set.

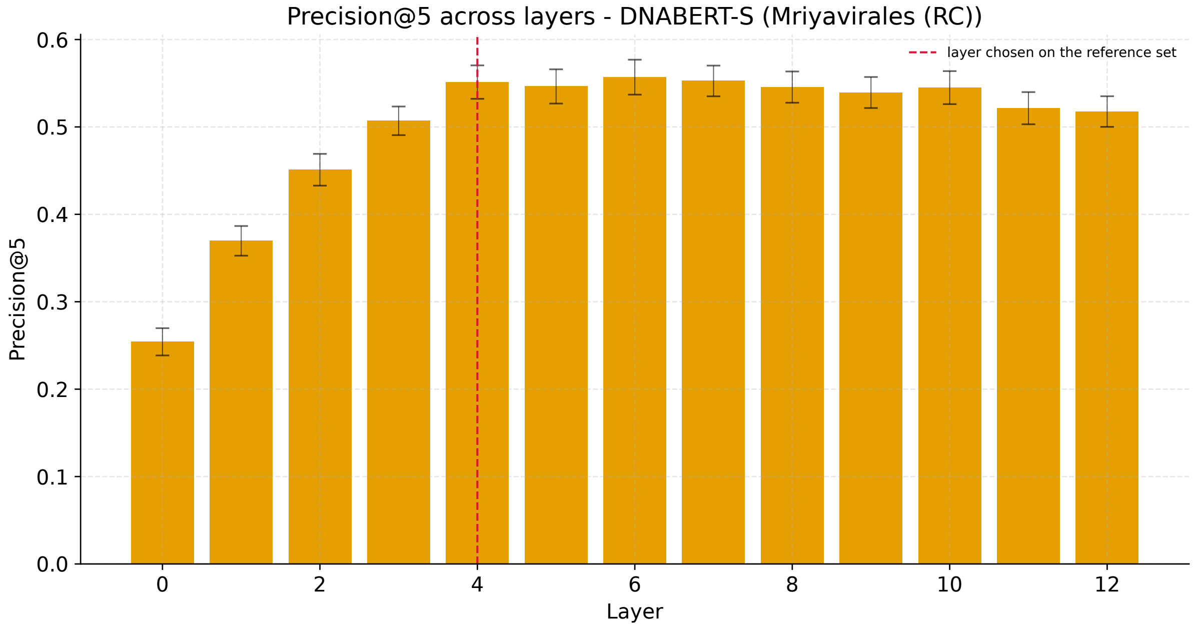


**Supplementary Figure S28.** Precision@5 on the reverse complements of mriyaviruses (RC) for each layer of DNABERT-S. Precision@5 was calculated using genome identity as the label, with the search space consisting of fragments of the corresponding genome and one reference subsample. Layers are numbered from 0 (embedding layer) to 12 (last transformer block). Bars show the mean ± standard deviation across the 50 reference subsamples. The dashed line marks the layer chosen on the reference set.

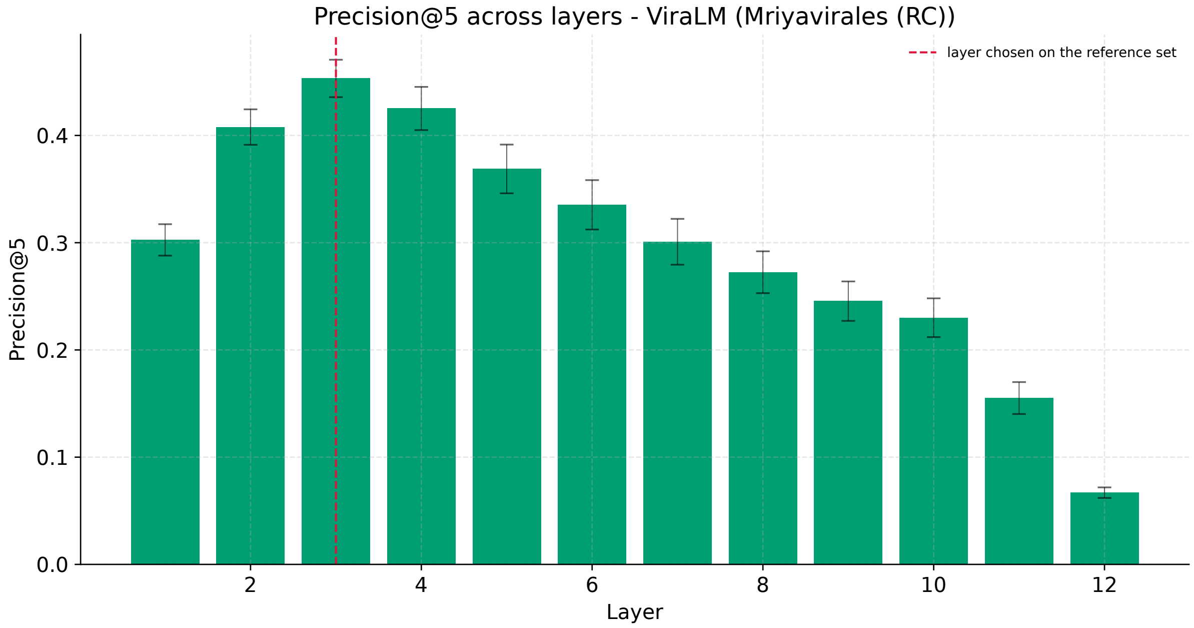


**Supplementary Figure S29.** Precision@5 on the reverse complements of mriyaviruses (RC) for each layer of ViraLM. Precision@5 was calculated using genome identity as the label, with the search space consisting of fragments of the corresponding genome and one reference subsample. Layers are the transformer blocks 1–12. Bars show the mean ± standard deviation across the 50 reference subsamples. The dashed line marks the layer chosen on the reference set.

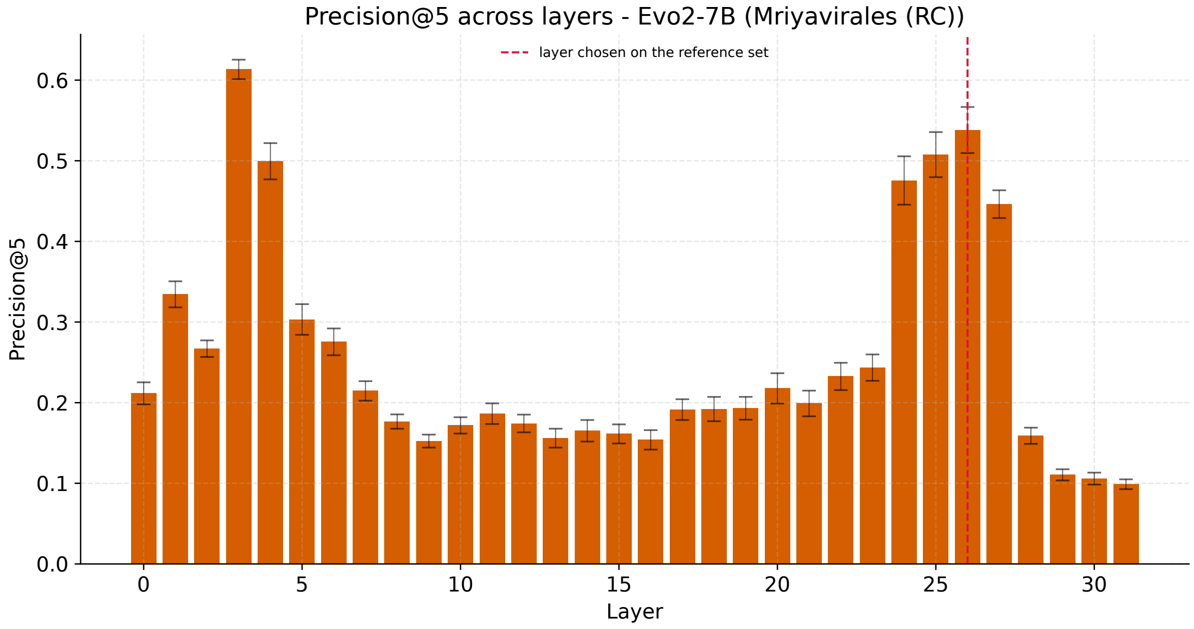


**Supplementary Figure S30.** Precision@5 on the reverse complements of mriyaviruses (RC) for each layer of Evo2-7B. Precision@5 was calculated using genome identity as the label, with the search space consisting of fragments of the corresponding genome and one reference subsample. Layers are the 32 blocks of the model, numbered 0–31. Bars show the mean ± standard deviation across the 50 reference subsamples. The dashed line marks the layer chosen on the reference set.

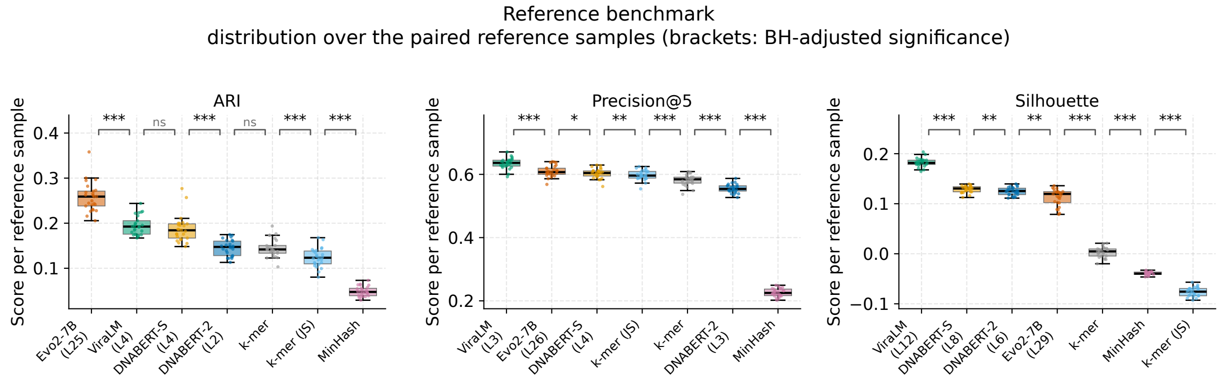


**Supplementary Figure S31.** Reference benchmark: distribution of ARI, precision@5 and the silhouette coefficient over the paired reference subsamples. For each method, the layer was chosen on the 25 even-numbered reference subsamples and the scores on the 25 odd-numbered subsamples are shown, so that the choice of the layer and the reported values share no subsample. Boxes show the median and quartiles, points show individual reference subsamples. Methods are ordered by their mean score; brackets between neighboring methods show Benjamini–Hochberg-adjusted p-values of two-sided paired Wilcoxon tests: * $p < 0.05$, ** $p < 0.01$, *** $p < 0.001$; ns, not significant.

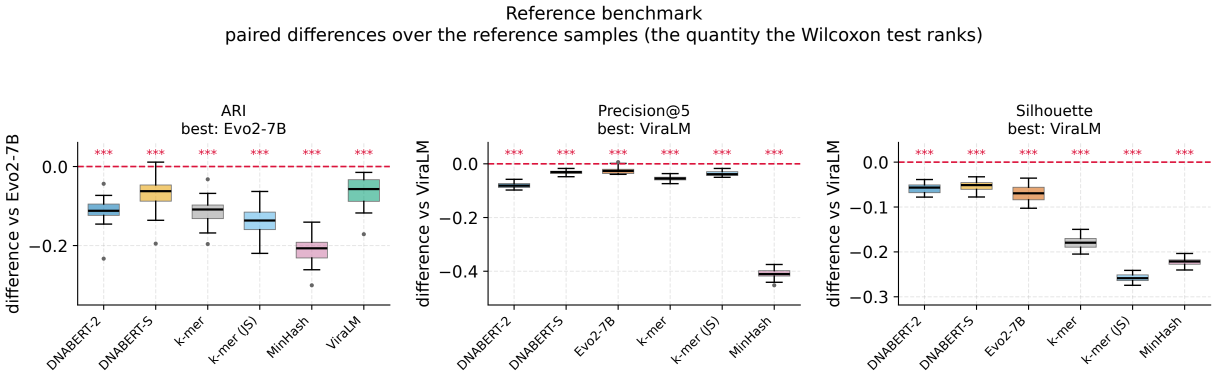


**Supplementary Figure S32.** Reference benchmark: ARI, precision@5 and the silhouette coefficient. Paired differences between each method and the best-scoring method of the panel (named above the panel), calculated for each reference subsample; this is the quantity ranked by the Wilcoxon test. Boxes show the median and quartiles, dots show outliers, the dashed line marks zero (no difference). For each method, the layer was chosen on the 25 even-numbered reference subsamples and the scores on the 25 odd-numbered subsamples are shown, so that the choice of the layer and the reported values share no subsample. Asterisks show Benjamini–Hochberg-adjusted p-values of two-sided paired Wilcoxon tests over the reference subsets: * $p < 0.05$, ** $p < 0.01$, *** $p < 0.001$; ns, not significant.

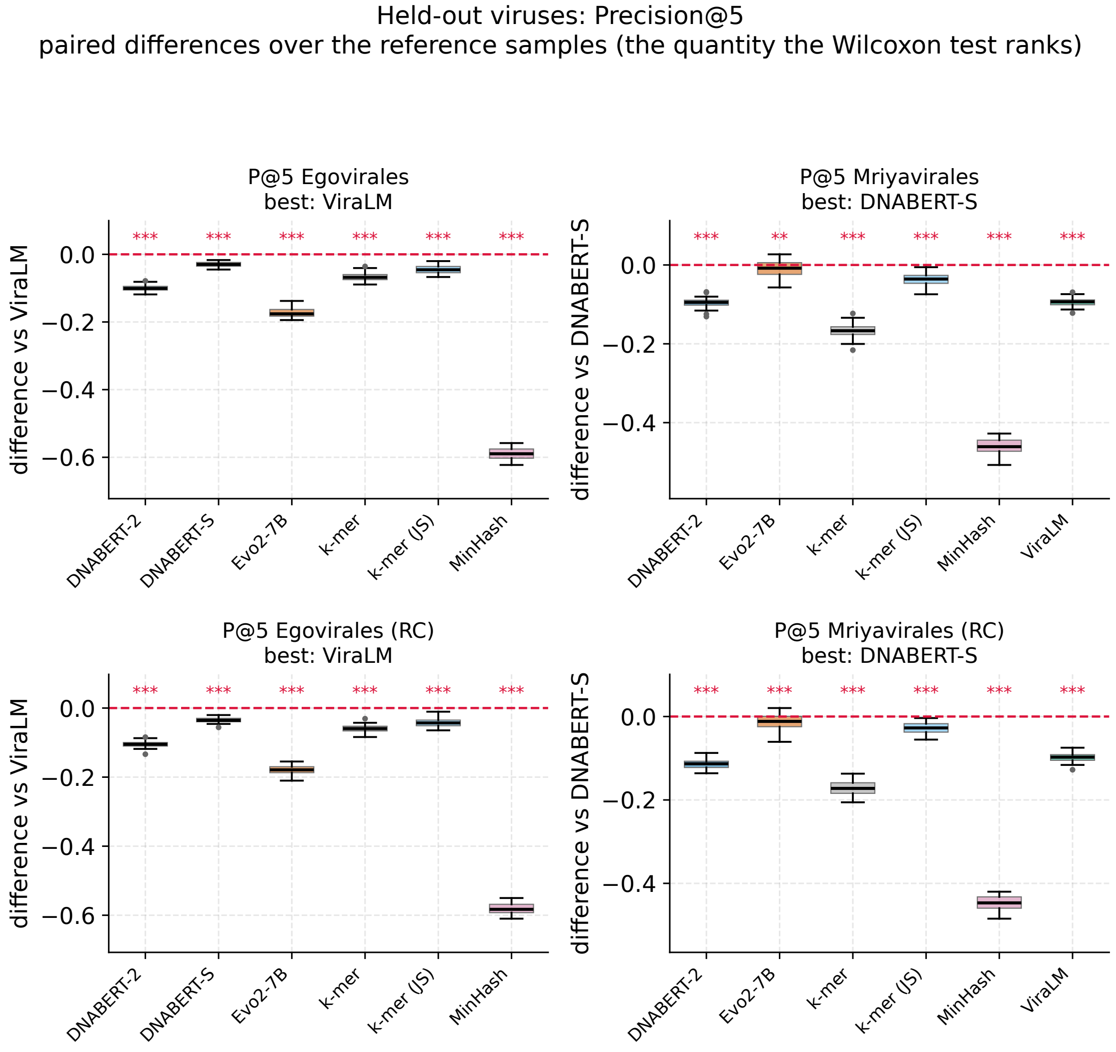


**Supplementary Figure S33.** Held-out viruses: precision@5 on egoviruses and mriyaviruses and their reverse complements (RC). Paired differences between each method and the best-scoring method of the panel (named above the panel), calculated for each reference subsample; this is the quantity ranked by the Wilcoxon test. Boxes show the median and quartiles, dots show outliers, the dashed line marks zero (no difference). The layer of each model is chosen on the reference set; the scores of all 50 reference subsamples are shown. Asterisks show Benjamini–Hochberg-adjusted p-values of two-sided paired Wilcoxon tests over the reference subsets: * $p < 0.05$, ** $p < 0.01$, *** $p < 0.001$; ns, not significant.

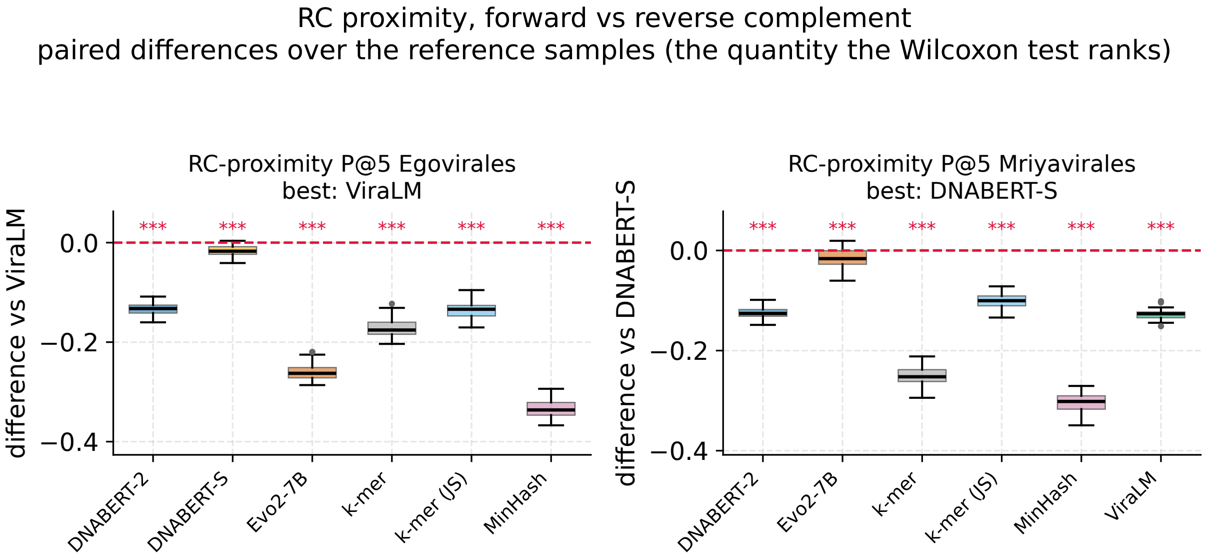


**Supplementary Figure S34.** Reverse-complement (RC) proximity (CROSS precision@5) for egoviruses and mriyaviruses. Paired differences between each method and the best-scoring method of the panel (named above the panel), calculated for each reference subsample; this is the quantity ranked by the Wilcoxon test. Boxes show the median and quartiles, dots show outliers, the dashed line marks zero (no difference). The layer of each model is chosen on the reference set; the scores of all 50 reference subsamples are shown. Asterisks show Benjamini–Hochberg-adjusted p-values of two-sided paired Wilcoxon tests over the reference subsets: * $p < 0.05$, ** $p < 0.01$, *** $p < 0.001$; ns, not significant.
